# Engineered coagulation factor VIII with enhanced secretion and coagulation potential for hemophilia A gene therapy

**DOI:** 10.1101/2024.12.05.626963

**Authors:** Yuji Kashiwakura, Yuto Nakajima, Kio Horinaka, Tiago J.S. Lopes, Yuma Furuta, Yuki Yamaguchi, Nemekhbayar Baatartsogt, Morisada Hayakawa, Yuko Katakai, Susumu Uchiyama, Osamu Nureki, Keiji Nogami, Tsukasa Ohmori

**Affiliations:** Department of Biochemistry, Jichi Medical University School of Medicine, 3311-1 Yakushiji, Shimotsuke, Tochigi 329-0498, Japan; Center for Gene Therapy Research, Jichi Medical University, 3311-1 Yakushiji, Shimotsuke, Tochigi 329-0498, Japan; Department of Pediatrics, Nara Medical University Hospital, 840 Shijo, Kashihara, Nara 634-8522, Japan; Department of Biological Sciences, Graduate School of Science, The University of Tokyo, Tokyo, Japan; Nezu Life Sciences, Heidelberg, Germany; Department of Biotechnology, Graduate School of Engineering, Osaka University, 2-1 Yamadaoka, Suita, Osaka 565-0871, Japan; U-Medico Inc., 2-1 Yamadaoka, Suita, Osaka 565-0871, Japan; The Corporation for Production and Research of Laboratory Primates, 1-16-2 Sakura, Tsukuba, Ibaraki 305-0003, Japan

## Abstract

The major challenges of gene therapy for hemophilia A using adeno-associated virus (AAV) vectors are reducing vector doses and the long-term maintenance of stable factor VIII (FVIII). Here, we developed engineered human B-domain-deleted FVIIIs (FVIIISQs) with enhanced secretion and coagulation potential. Intracellular accumulation was markedly reduced in some engineered FVIIISQs, resulting in reduced unfolded protein responses. The administration of AAV vectors carrying engineered FVIIISQ to hemophilia A mice resulted in approximately eight-fold higher FVIII activity and four-fold higher FVIII antigen levels compared with wild-type FVIIISQ administration. The specific FVIII activity of the engineered FVIIISQ was 3.6 times higher than that of the wild-type FVIIISQ, and its binding to activated coagulation factor IX was significantly enhanced, which is supported by the structural analysis. In macaques, the administration of AAV5 vector carrying the engineered FVIIISQ without CpG sequences resulted in a supra-physiological increase in plasma FVIII activity at a dose one-thirtieth that of valoctocogene roxaparvovec (2 × 10^12^ vg/kg). The engineered FVIIISQ may thus provide stable, long-term therapeutic efficacy in AAV-mediated hemophilia A gene therapy even at low doses.

---

Hemophilia A is an X-linked recessive bleeding disorder that is caused by an abnormality in the *F8* gene. Current treatment of hemophilia A primarily involves replacing the deficient factor VIII (FVIII) protein in the patient’s blood. Although the recent developments of bispecific antibodies and other therapies have greatly improved patients’ quality of life, lifelong drug administration remains necessary. Gene therapy using adeno-associated virus (AAV) vectors may address this issue by providing sustained therapeutic levels of coagulation factors. In clinical trials, the intravenous administration of AAV vectors carrying FVIII to hemophilia A patients significantly increases plasma FVIII activity into the therapeutic range without serious adverse events^1–5^. However, long-term follow-up data indicate a gradual decrease in FVIII expression levels over time, suggesting that therapeutic effects may not be lifelong^6^.

Gene therapy for hemophilia using AAV vectors targets liver hepatocytes for the expression of therapeutic genes. Because *F8* is expressed in liver sinusoidal endothelial cells^7^, gene therapy for hemophilia A involves expressing the target gene in cells that do not physiologically produce it. It has been suggested that the ectopic expression of the very large FVIII protein in hepatocytes causes an unfolded protein response (UPR)^8,9^, which may lead to reduced gene expression over time^6^. Although UPR functions as an adaptive signaling pathway that prevents misfolded proteins in the endoplasmic reticulum (ER), it can lead to cell death via apoptosis as well as translational arrest and protein degradation, which reduce the protein load in the ER^10^. The immunohistochemical analysis of liver specimens from hemophilia A patients collected 2.6–4.1 years after gene therapy revealed no evidence of UPR^11^. However, there is not always a correlation between mRNA and protein expression levels, and patients with lower mRNA levels can achieve relatively stable FVIII expression^11^. Further, inter-individual differences in FVIII expression may be influenced by transduction efficacy as well as unknown host-mediated post-translational mechanisms^11^. These data indicate that preventing FVIII accumulation within cells after its ectopic translation may lead to more stable therapeutic effects.

Another issue with gene therapy for hemophilia A is the requirement for high doses of AAV vectors^6^, although the blood molar concentration of FVIII is lower than that of factor IX (FIX)^12^. High doses of AAV vectors reportedly cause immune-related adverse events such as liver dysfunction and thrombotic microangiopathy^13^. High doses of vectors also elicit an immune response, thus increasing the need for immunosuppressive agents in AAV vector-mediated gene therapy^14^. The prevention of FVIII accumulation within cells may also facilitate increased plasma levels of FVIII, thereby reducing the required vector doses. Another way to further reduce the vector dosage requirement is to enhance coagulation factor function. Indeed, effective gene therapy at low doses has been achieved for hemophilia B using FIX Padua^15^. FIX Padua features an R338L substitution (identified from a family with thrombophilia) that results in activity that is eight times that of wild-type FIX^16^.

To resolve these issues for hemophilia A gene therapy, several attempts at FVIII protein engineering, including for improved protein stability^17,18^, single chain formation^19,20^, enhanced secretion^8,21^, and increased expression^4,22^, have been reported. However, despite these efforts, insufficient preclinical data have demonstrated their effectiveness in clinical practice at low vector doses. Here, we aimed to develop an engineered FVIII with enhanced activity and secretion, to allow for human hemophilia A gene therapy with stable gene expression using a low vector dose.

## Results

### FVIII engineering based on non-human mammal FVIII

FVIII activity (FVIII:C) varies between animal species^22–25^. Canine and porcine FVIII reportedly have higher FVIII:C than human FVIII^22,25^. We therefore compared the FVIII:C of several mammals. Human amino acid sequences in B-domain-deleted FVIIIs (FVIIISQs) were relatively conserved among canine, porcine, bovine, and ovine species, differing from human FVIIISQ by just 13%–16% (Supplementary Fig. 1a,b). FVIII:C in supernatant from HEK293 and Huh-7 (a human hepatocyte cell line) cells after transient transfection was higher with FVIIISQs derived from non-human mammals (Supplementary Fig. 1c,d).

We next constructed four types of engineered FVIIISQs based on the amino acid sequences of these animal species, as follows (Supplementary Table 1 and Supplementary Fig. 2a):

FVIIISQ(Ver.1): substitution of 72 amino acids common to canine and porcine FVIII.

FVIIISQ(Ver.2): substitution of 58 amino acids common to canine, porcine, bovine, and ovine FVIII.

FVIIISQ(Ver.3): the 21 substituted amino acids in FVIIISQ(Ver.1) that had unchanged physiochemical properties (e.g., hydrophilic and negatively charged, D and E; hydrophilic, S and T; hydrophilic and positively charged, K, H, and R; hydrophobic and aliphatic, I, L, and V) were restored to the original human sequences (51 amino acid substitutions remained).

FVIIISQ(Ver.4): substitution of the 36 amino acids common to FVIIISQ(Ver.2) and FVIIISQ(Ver.3).

The FVIII:C of the engineered FVIIISQs were significantly higher in the supernatant of Huh-7 cells after transient transfection (Supplementary Fig. 2b). These activities were much higher than those of previously reported engineered FVIII (namely, ET3, X5, and JF12) (Supplementary Fig. 2b)^21,26,27^. We further generated AAV8 vectors harboring FVIIISQs and intravenously administered them into hemophilia A model mice (Fig. 1a). The AAV vector-mediated expression of engineered FVIIISQ resulted in significantly higher plasma FVIII:C (Fig. 1b,c). The most effective version was FVIIISQ(Ver.4), which showed 8.2-fold higher plasma FVIII:C than wild-type FVIIISQ in the one-stage assay and 3.4-fold higher FVIII:C in the chromogenic assay (Fig. 1b,c). The increases in FVIII:C by AAV harboring FVIIISQ(Ver.4) were also higher than those by AAV harboring X5 (Fig. 1b,c). Plasma FVIII antigen (FVIII:Ag) levels were also increased four-fold in the engineered FVIIISQ(Ver.4) (Fig. 1d) compared with wild-type FVIIISQ. However, AAV vector genome (vg) and FVIII mRNA levels in the liver were equivalent or reduced compared with wild-type FVIIISQ (Fig. 1e,f).

**Fig. 1.**
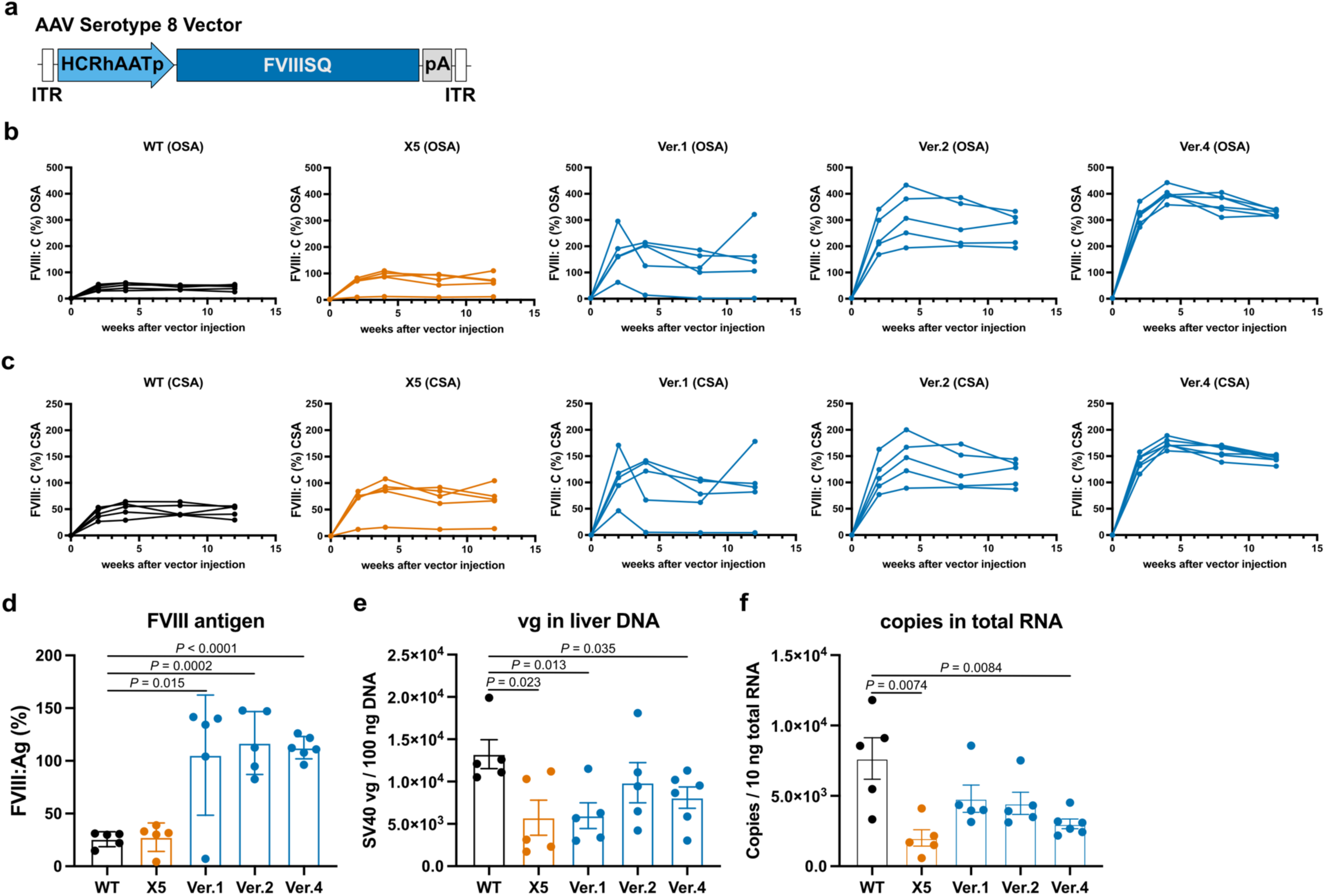
Expression of FVIII by the AAV8 vector harboring FVIIISQs in hemophilia A model mice. **(a)** Structure of the AAV vector. **(b–f)** AAV8 vectors were intravenously administered into male hemophilia A model mice (1 × 10^11^ vg/kg). **(b, c)** Plasma FVIII activity (FVIII:C) assessed using the one-stage clotting assay (**b**, OSA) or chromogenic assay (**c**, CSA). **(d)** FVIII antigen (FVIII:Ag) at 4 weeks after vector injection. **(e)** AAV vector copy number in liver tissue. **(f)** *F8* mRNA expression in liver (*n* = 5–6 in each experiment). WT: wild-type FVIII, X5: engineered FVIII with five amino acid substitutions from a previous study, Ver: engineered FVIII in our study. Values are presented as the mean ± SEM (WT, X5, Ver.1, and Ver.2, *n* = 5; Ver.4, *n* = 6). Significance was assessed using Student’s *t*-tests.

### UPR induction and intracellular FVIII accumulation

Next, we assessed UPR induction upon FVIII expression *in vitro*. We established Huh-7 cells that expressed luciferase upon the splicing of X-box binding protein 1 by the activation of inositol-requiring enzyme 1 (IRE1), a proximal sensor of UPR (Fig. 2a)^28^. The expression of wild-type FVIIISQ induced a significant UPR response, whereas FVIIISQ(Ver.2) and FVIIISQ(Ver.4) clearly showed less UPR induction (Fig. 2b). We further assessed intracellular FVIII accumulation by examining protein levels via immunoblotting of cell lysates and supernatants of Huh-7 cells transfected with each FVIIISQ. The engineered FVIIISQs had higher protein levels in the supernatant compared with wild-type FVIIISQ (Fig. 2c,d), and the protein levels of FVIIISQ(Ver.2) and FVIII SQ(Ver.4) resulted in a significantly lower retention of light chains within the cells (Fig. 2e,f).

**Fig. 2.**
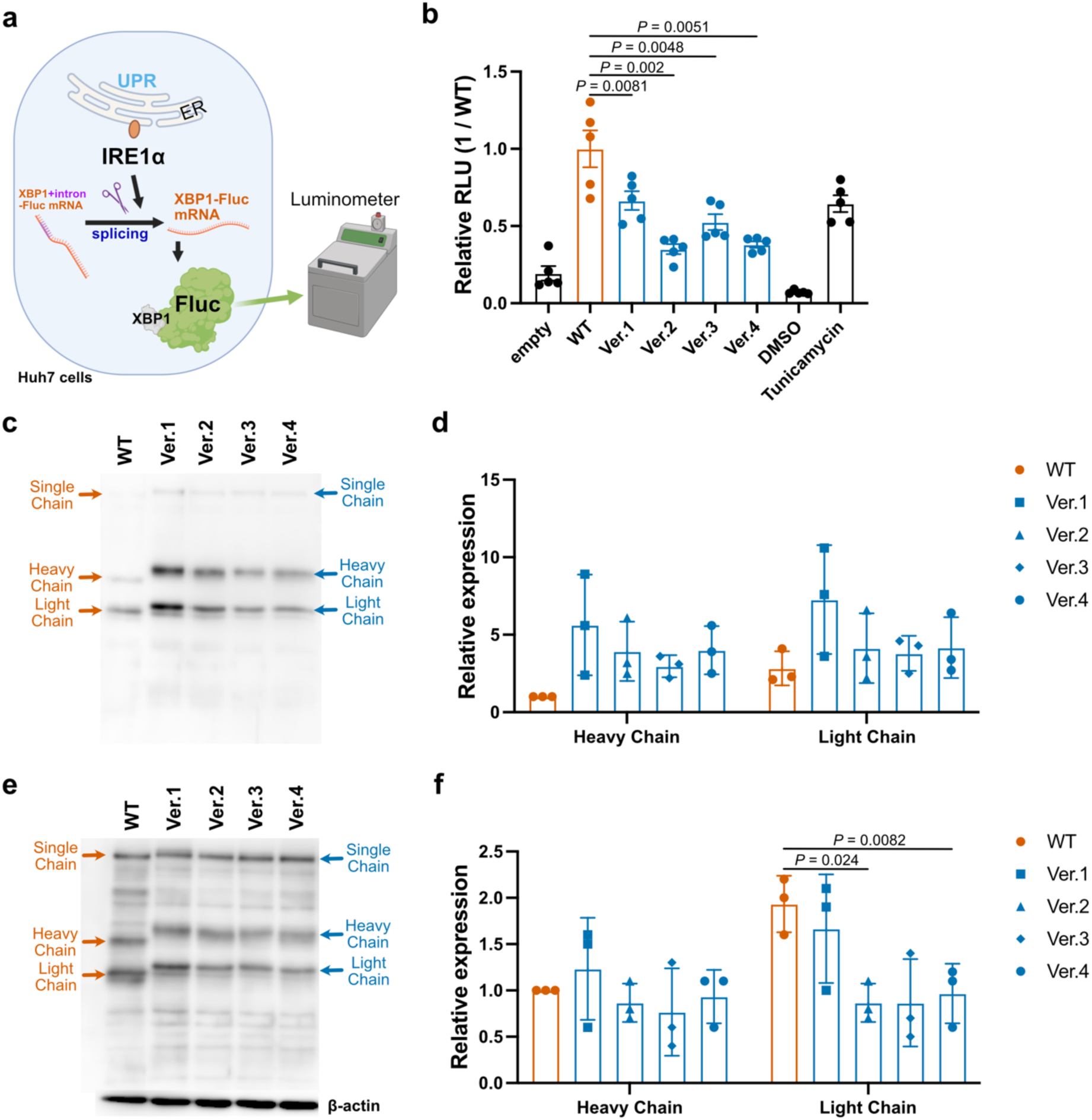
Unfolded protein response and intracellular accumulation of FVIII. **(a, b)** We established the Huh-7 cells to analyze the unfolded protein response using luciferase expression. **(b)** Luciferase expression of the cells after plasmid transduction. Values are presented as the mean ± SEM (*n* = 5). **(c–f)** Huh-7 cells were transduced with plasmids expressing the FVIIISQs. FVIII protein expression in the supernatant **(c,d)** and cell lysates **(e,f)**. WT: wild-type FVIII, Ver: engineered FVIII in our study. Values are presented as the mean ± SEM (*n* = 3). Significance was assessed using Student’s *t*-tests.

The greater molecular weight of the heavy and light chains of engineered FVIIISQs compared with those of wild-type FVIIISQs suggested the existence of additional post-translational modifications of engineered FVIIISQs (Fig. 2). To analyze the amino acid substitution responsible for greater molecular weights, we reverted each FVIIISQ(Ver.4) domain into the corresponding human domain (Supplementary Fig. 3). The substitution of K213N made the heavy chain contain NNS, a typical N-X-T/S motif that is usually *N-*glycosylated^29^. As expected, we detected the *N-*glycosylation of N213 in FVIIISQ(Ver.4), with clear fragment ions corresponding to the *N-*glycan core structure (Supplementary Fig. 4). In addition, although no post-translational modifications, such as hydroxylation, of proline in P1657 were detected, *O-*glycosylation was observed at T1653 and/or T1654 near P1657 (Supplementary Fig. 4).

### Biochemical and structural analysis of FVIII

FVIII acts as a cofactor for the proteolytic cleavage of factor X (FX) substrate by activated FIX enzyme (FIXa). The binding of activated FVIII (FVIIIa) to FIXa on activated platelet surfaces forms the intrinsic “tenase” complex, which catalyzes FX to its activated form (FXa). FVIIIa is easily inactivated by the dissociation of the A2 domain after activation because of the non-covalent weak electrostatic interactions between the A1 and A2 domains in FVIII^30^. To characterize the biochemical properties of FVIIISQ(Ver.4) protein, we purified FVIIISQ(Ver.4) and wild-type FVIIISQ proteins from the supernatants of HEK293 cells stably expressing each protein. FVIIISQ(Ver.4) showed 3.6- and 1.5-fold higher FVIII:C than wild-type FVIIISQ in the one-stage and chromogenic assays, respectively (Supplementary Table 2). The kinetic parameters are summarized in Fig. 3a,b. The *K*_cat_/*K*_m_ of FXa generation in FVIIISQ(Ver.4) for FX appeared to be comparable with that of wild-type FVIIISQ (Fig. 3a). By contrast, the apparent *K*_d_ for FIXa in FVIIISQ(Ver.4) (0.57 ± 0.07 nM) was significantly lower than that in wild-type FVIIISQ (1.3 ± 0.2 nM) (Fig. 3b). In addition, the FXa generated in FVIIISQ(Ver.4) (85 ± 6.7 nM FXa/min/nM FVIII) was significantly greater than that in wild-type FVIIISQ (46 ± 2.0 nM FXa/min/nM FVIII). These data indicate that the interaction between FIXa and FVIIISQ(Ver.4) in the tenase complex is enhanced compared with that of wild-type FVIIISQ. Moreover, the thrombin generation potential of FVIIISQ(Ver.4) was equivalent to that of wild-type FVIIISQ (Fig. 3c). The spontaneous decay of FVIIISQ(Ver.4) was assessed by measuring FVIII activity after activation by thrombin using a one-stage clotting assay. The decline in FVIIIa activity in FVIIISQ(Ver.4) (0.055 min^−1^) was approximately 2.6-fold faster than in wild-type FVIIISQ (Fig. 3d), indicating the rapid dissociation of the A2 domain.

**Fig. 3.**
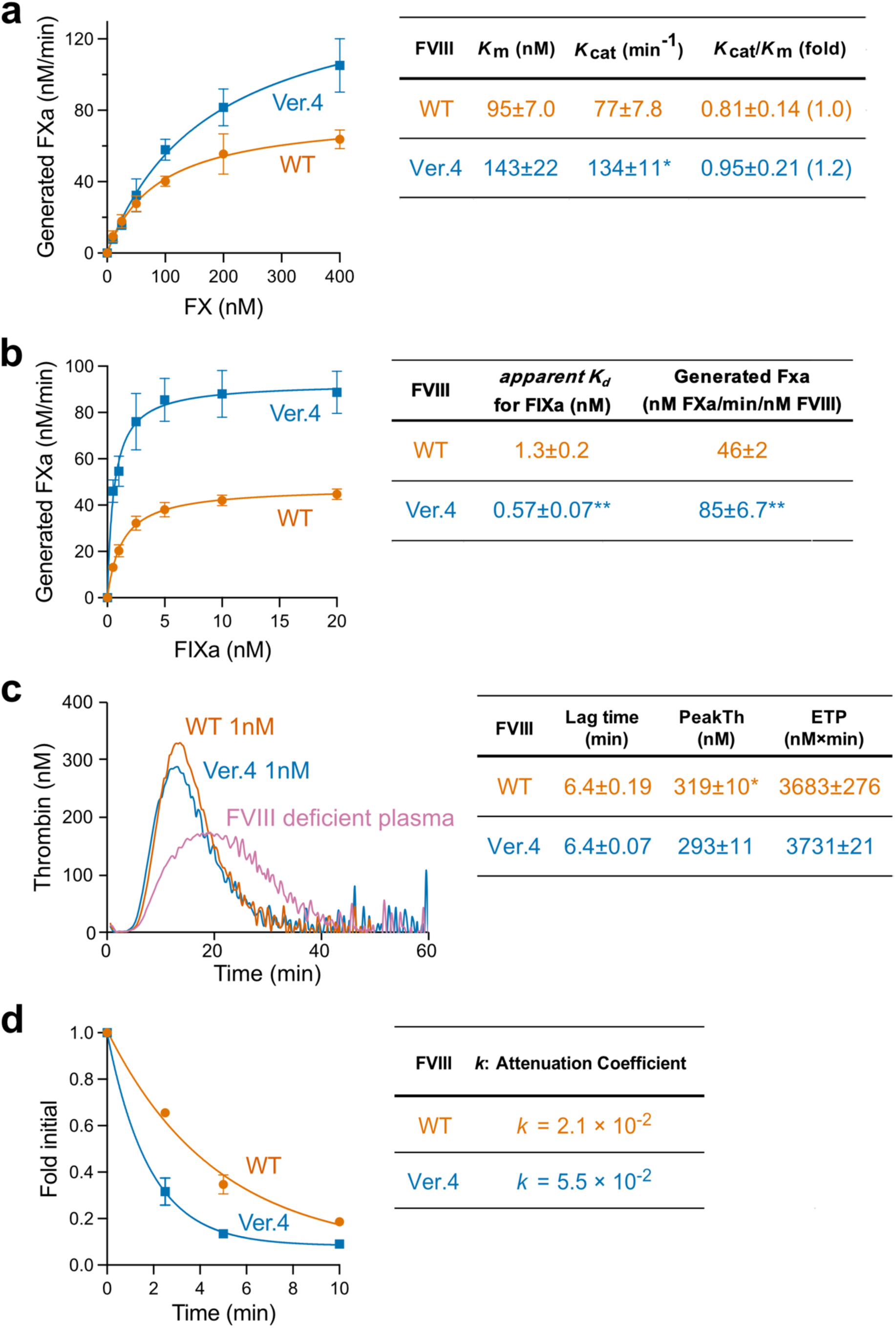
Biochemical analysis of FVIII protein. **(a, b)** FVIII (1 nM) was activated by thrombin (30 nM) for 30 s in the presence of phospholipid vesicles (20 μM). **(a)** Association with FX. FXa generation was initiated by the addition of FIXa (40 nM) and various concentrations of FX. **(b)** Association with activated FIX. FXa generation was initiated by the addition of FX (300 nM) and various concentrations of FIXa. **(c)** A2 dissociation after thrombin activation. FVIII (1 nM) was incubated with thrombin (30 nM) prior to measuring FVIII activity at the indicated times using a one-stage clotting assay. **(d)** Thrombin generation potential. FVIII (1 nM) was added to FVIII-deficient plasma. These samples were then incubated with recombinant tissue factor (1 pM) and phospholipid vesicles (4 µM) prior to the addition of fluorogenic substrate and CaCl_2_ at the start of the assay. WT: wild type FVIIISQ, Ver.4: FVIIISQ(Ver. 4).

To obtain structural insights into the unique biochemical properties observed in FVIIISQ(Ver.4), we resolved the cryo-electron microscopy structure of purified FVIIISQ(Ver.4) protein at 3.17 Å (Fig. 4a). The structure revealed that the purified FVIIISQ(Ver.4) protein consisted of A1–3 and C1–2 domains (Fig. 4b). By superimposing the structure of FVIIISQ(Ver.4) protein onto that of a previously reported wild-type FVIII protein (model modified from PDB:6MF2, see Supplementary Fig. 5 for details), we demonstrated that, except for the C2 domain, the overall architecture was highly conserved (Cα root mean square deviation = 0.848 Å); however, multiple conformations were observed in the C2 domain of FVIIISQ(Ver.4) protein, suggesting its intrinsic flexibility (Fig. 4c).

**Fig. 4.**
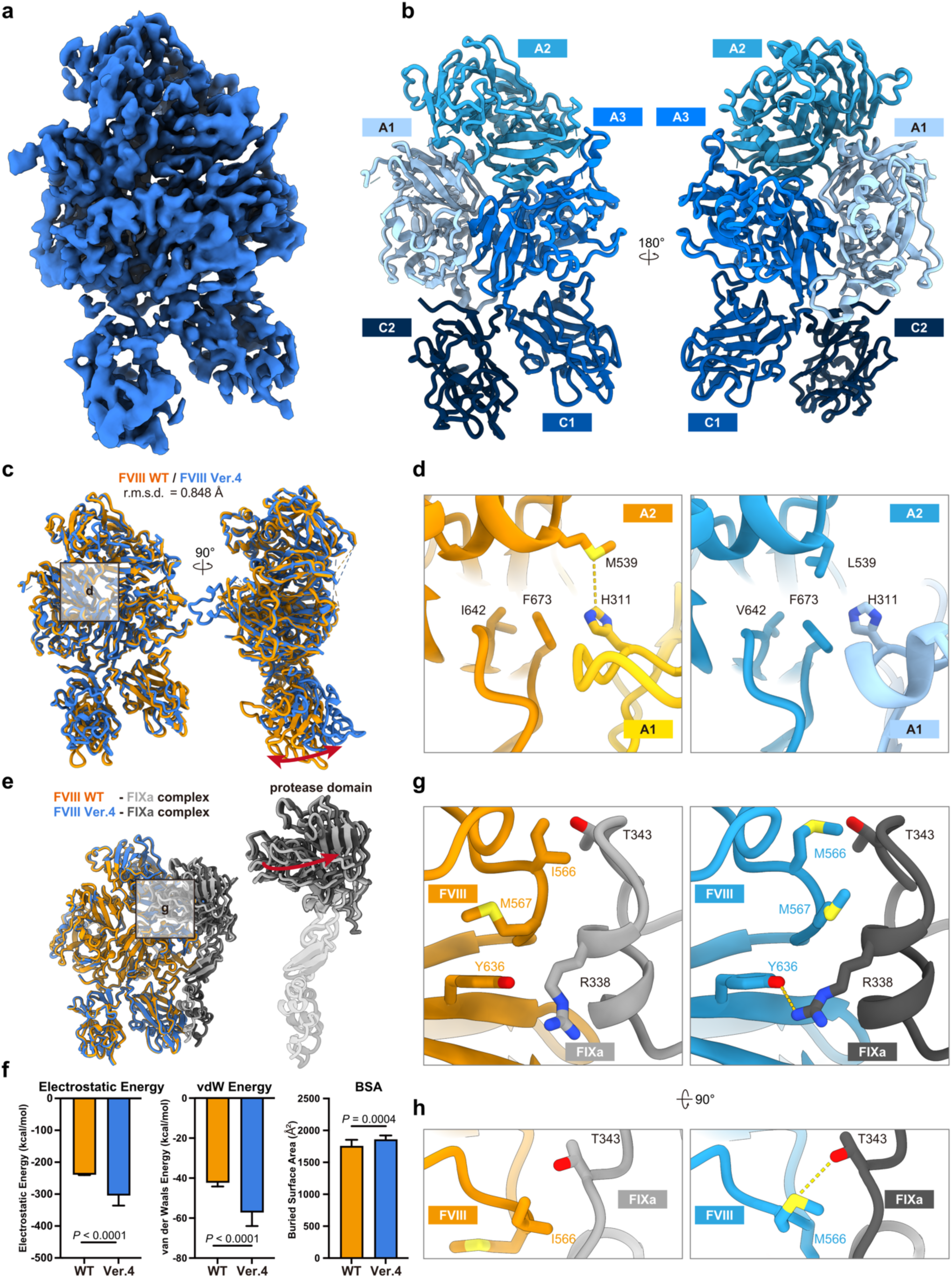
Structural analysis of the engineered FVIII. **(a, b)** Cryo-electron microscopy map (a) and structural model (b) of FVIIISQ(Ver.4). **(c)** FVIIISQ(Ver.4) (blue) superimposed onto wild-type FVIIISQ (orange). The conformational shift of the C2 domain is depicted by the red arrow. **(d)** The A2–A1 interface of wild-type FVIIISQ (left) and FVIIISQ(Ver.4) (right). The yellow dashed line represents the interaction between M539 and H311. The distance is 5.0Å. **(e)** FVIIISQ(Ver.4) (blue)–FIXa (dark gray) complex superimposed onto a wild-type FVIIISQ (orange)–FIXa (light gray) complex. The shift in the protease domain of FIXa is depicted by the red arrow. **(f)** Results of the HADDOCK simulation. Values are presented as the mean ± standard deviation (WT, *n* = 20; Ver.4, *n* = 28). Significance was assessed using Welch’s *t*-tests. **(g, h)** Side **(g)** and top **(h)** views of the FVIII–FIXa interface with wild-type FVIIISQ (left) and FVIIISQ(Ver.4) (right). The yellow dashed lines represent the Y636–R338 and M566–T343 hydrogen bonds. The distances are 3.2 and 4.2 Å, respectively.

To explore the structural basis for the fast-inactivating nature of FVIIISQ(Ver.4), we next focused on the interface between A2 and A1/A3. Although the interface between the A2 and A3 domains was similar to that of wild-type FVIII, A2 interacted differently with A1 in FVIIISQ(Ver.4). In wild-type FVIII, M539 in the A2 domain interacts with H311 in the A1 domain, forming a so-called Met-aromatic motif^31^. The conformation of H311 in wild-type FVIII is supported by the stacking interaction with F673, whose conformation is defined synergistically by a steric strain posed by surrounding residues, including I642. In FVIIISQ(Ver.4), however, the M539L substitution eliminates this interaction, thereby enabling the rapid dissociation of the A2 subunit from the rest (Fig. 4d).

To investigate the potential influence of introduced substitutions on FIXa binding, we predicted the structure of both wild-type FVIIISQ and FVIIISQ(Ver.4) in complex with FIXa using AlphaFold 3. Both complex structures shared a similar binding mode of FIXa protease domain onto the FVIII A2 domain, as previously reported^32^. Nonetheless, although the predicted structure of the wild-type FVIIISQ– FIXa complex superposed onto that of the FVIIISQ(Ver.4)–FIXa complex very well, the protease domain in the latter case was slightly shifted away from the FVIII A2 domain compared with the former case (Fig. 4e). To quantify the difference in binding modes, we performed docking simulations using HADDOCK 2.4. The result indicated that electrostatic energy, van der Waals energy, and buried surface area were preferable in the case of FVIIISQ(Ver.4), indicating stronger and more extensive interactions with FIXa compared with wild-type FVIIISQ (Fig. 4f). In fact, the predicted structures showed that M567 was angled toward FIXa in the case of FVIIISQ(Ver.4), thus providing a van der Waals interaction between FVIII and FIXa. This conformational difference seems to be influenced by the I566M substitution, with the deletion of isoleucine’s methyl group on its β-carbon. This M567 conformation also likely repels the helix of FIXa, providing space for R338 in FIXa to extend and form a hydrogen bond with Y636 in FVIII (Fig. 4g). Furthermore, M566 is located relatively close to T343 in FIXa and is speculated to form a hydrogen bond with it, resulting in decreased electrostatic energy (Fig. 4h). Overall, these observations might explain why FVIIISQ(Ver.4) can bind FIXa more strongly than wild-type FVIIISQ.

### Immunogenicity of FVIII

One of the greatest challenges in hemophilia A patient care is the development of neutralizing antibodies (FVIII inhibitors) against the FVIII protein that is used for replacement therapy. To assess the risk of inhibitor induction as a result of amino acid substitutions, the immunogenicity of FVIIISQ(Ver.4) was assessed using *in silico* analysis. We used NetMHCIIpan^33^ to calculate and compare the affinities of the epitopes present in wild-type FVIIISQ and FVIIISQ(Ver. 4) against the 46 most common human leukocyte antigen (HLA) types in humans (Fig. 5, Supplementary Table 3). Together, these HLA types should cover approximately 90% of the population^34^. Additionally, to quantify the ‘risk’ of an immune response in people harboring specific HLA types, we developed a measure called the T-Score, which was defined as the allele frequency divided by the affinity concentration (Fig. 5, Supplementary Table 3). In this T-Score, higher values indicate a higher affinity of an epitope to a common HLA type in the population, which might lead to T-cell activation and subsequent inhibitor development^35^. We observed that wild-type FVIIISQ and FVIIISQ(Ver. 4) shared many epitopes (Fig. 5, Supplementary Table 3); however, because these epitopes do not lead to an immune response in the general population, they are likely to be tolerated by the immune system^36^. Moreover, we observed the presence of another 16 novel combinations of FVIIISQ(Ver.4) epitopes and HLA types, but these epitopes had low T-Scores (<0.05), suggesting that they do not bind to the most common HLAs with high affinity (Supplementary Tables 4 and 5). Together, these results suggest that the vast majority of epitopes in the novel engineered FVIIISQ(Ver.4) are comparable with those in its wild-type version; although some epitopes were introduced, they likely bind with low affinity to specific HLA types (Supplementary Table 5).

**Fig. 5.**
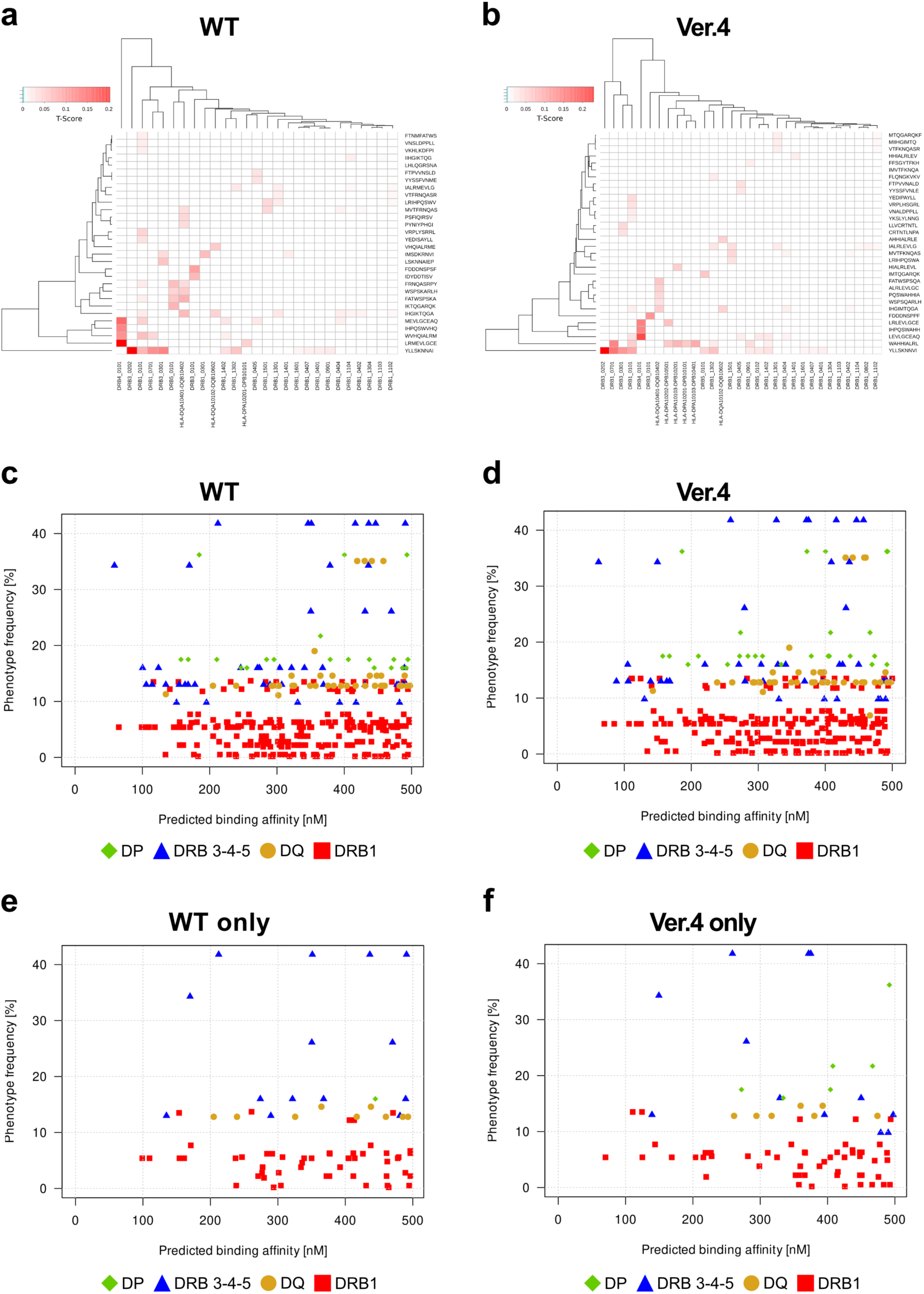
*In silico* assessment of immunogenicity. The predicted binding affinities of the FVIIISQ WT and Ver.4 epitopes to the most common HLA subtypes in the human population are depicted. **(a,b)** T-Score, defined as the allele frequency / affinity concentration. Higher values indicate a stronger affinity of an epitope to an HLA type that is relatively frequent in the population. Although the engineered FVIIISQ Ver.4 had novel epitopes, the vast majority had small T-Score values. **(c,d)** Binding affinities of epitopes against the *DRB1*, *DRB3*, *DRB4, DRB5*, *DP*, and *DQ* HLA types. Only epitopes with a binding affinity < 500nM are shown, and the proteins were predicted to have comparable binding profiles. **(e,f)** Binding of epitopes exclusive to either the wild-type or engineered versions of the FVIIISQ protein. Again, the proteins had similar profiles, and very few epitopes were predicted to strongly interact with the most frequent HLA types in the population.

### Non-human primate trial

Finally, we conducted a preclinical study in non-human primates; specifically, in macaques (*Macaca fascicularis*). A previous clinical trial using AAV vectors suggested that the innate immunity induced by CpG sequences in the transgene results in a loss of transgene expression in humans^37^. The FVIIISQ cDNA sequence used in previous experiments has already been codon-optimized^38^, but contained more CpG sequences after the optimization. We therefore designed several codon-optimized FVIIISQ(Ver. 4) sequences without CpG sequences for the macaque experiments. The mouse transthyretin (mTTR) promoter has less CpG sequences than the HCRhAAT chimeric promoter (an enhancer element of the hepatic control region of the Apo E/C1 gene and the human anti-trypsin promoter). We therefore removed all CpG sequences from the mTTR promoter, which did not affect promoter activity (Supplementary Fig. 6a,b). Next, we designed a plasmid harboring codon-optimized FVIIISQ(Ver. 4) driven by the mTTR promoter without any CpG sequences, and obtained a significantly higher FVIII:C than with the original FVIIISQ(Ver. 4) *in vitro* (Supplementary Fig. 6c,d). Administration of the AAV8 vector in hemophilia A model mice resulted in similar findings (Supplementary Fig. 6c,e).

We further administered AAV8 and AAV5 vectors harboring the codon-optimized FVIIISQ(Ver.4) sequence without CpG into macaques (AAV8, 2 × 10^12^ vg/kg [*n* = 2]; AAV5, 6 × 10^12^ vg/kg [*n* = 2] and 2 × 10^12^ vg/kg [*n* = 3]) (Fig. 6). The peak increase in plasma FVIII:C after administration far exceeded the human reference values (Fig. 6a,b). Human FVIII:Ag, assessed using human FVIII-specific enzyme-linked immunosorbent assay, was also significantly increased after administration (AAV8 [2 × 10^12^ vg/kg], 50.46% and 186.42% [*n* = 2]; AAV5 [6 × 10^12^ vg/kg], 67.78% and 104.98% [*n* = 2]; AAV5 [2 × 10^12^ vg/kg], 19.92%, 62.07%, and 63.21% [*n* = 3]) (Fig. 6c). The plasma FVIII levels then gradually decreased in six of the seven macaques as a result of the emergence of neutralizing antibodies against human FVIII (Supplementary Fig. 7). We did not detect any significant changes in laboratory data, including alanine amino transferase (Supplementary Fig. 8).

**Fig. 6.**
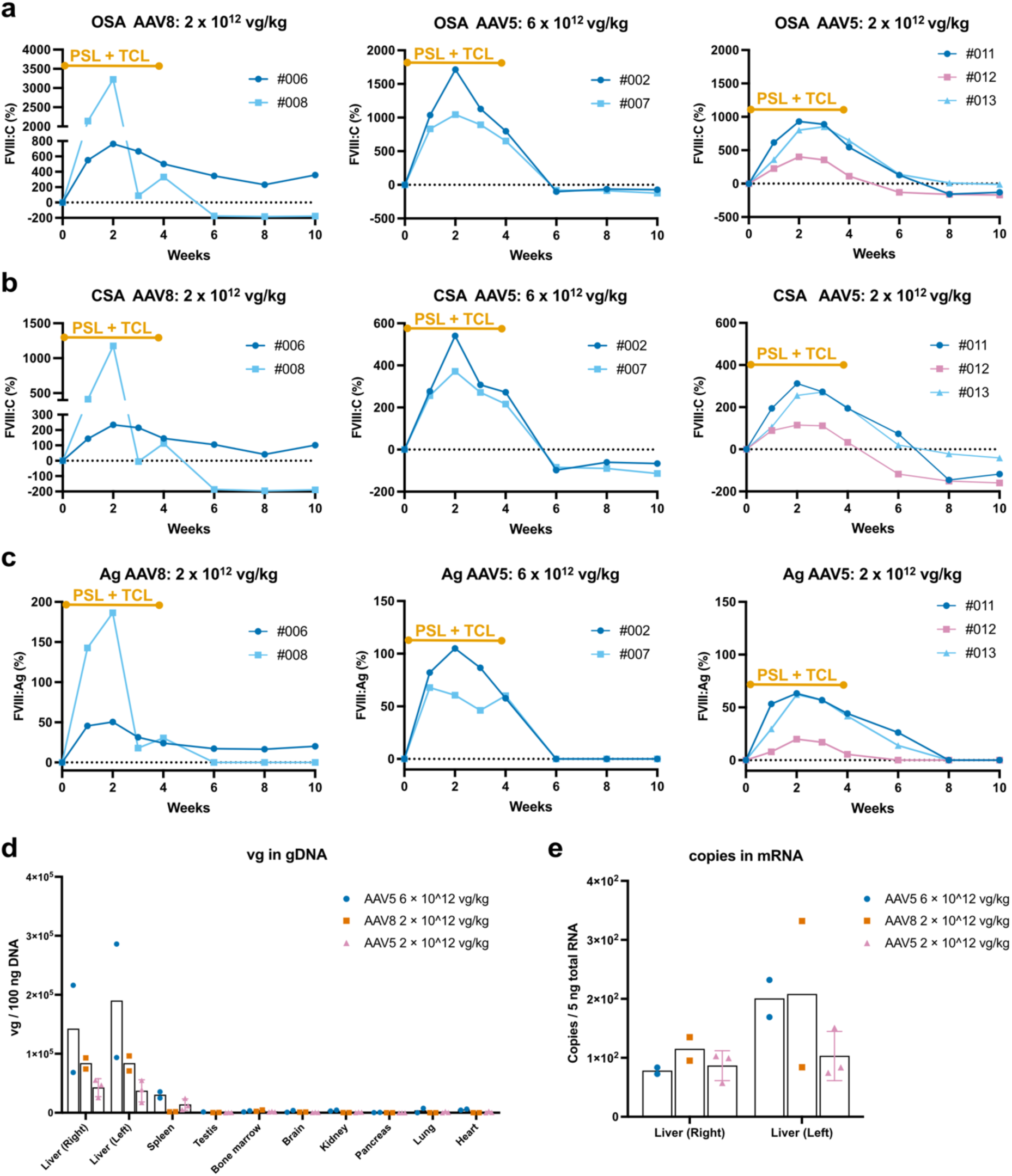
Expression of FVIII by AAV vector harboring the engineered FVIIISQ in macaques. **(a–c)** AAV8 or AAV5 harboring the codon-optimized engineered FVIIISQ(Ver.4) driven by the mouse transthyretin promoter without CpG sequences was administered to male macaques through a peripheral vein (AAV8: 2 × 10 vg/kg^12^, #6, #008; AAV5: 6 × 10^12^ vg/kg, #002, #007; AAV5: 2 × 10^12^ vg/kg, #011, #012, #013). The increase in plasma FVIII activity (FVIII:C) was assessed using the one-stage clotting assay (OSA) **(a)** or chromogenic assay (CSA) **(b)**, and the FVIII antigen (FVIII:Ag) was measured by ELISA **(c). (d)** AAV genome in an indicated organ. **(e)** mRNA expression in the liver.

After these observations, we obtained specimens to analyze the vector genome in organs. In the liver, AAV vector genome and FVIIISQ(Ver.4) mRNA expression were preserved even in macaques with reduced plasma FVIII (Fig. 6d,e), indicating the maintenance of the transduced cells. AAV genomes were predominantly present in the liver, and there was a higher tendency for vector distribution to the spleen with AAV5 than with AAV8 (Fig. 6d). We further quantified the distribution of AAV vectors in detail using digital polymerase chain reaction (PCR). Although 0.16–0.85 AAV vector copies were detected in haploid genome of the liver, only marginal AAV genome was detected in the testes (Supplementary Fig. 9).

## Discussion

Gene therapy using AAV vectors is a breakthrough modality that enables the long-term cure of hemophilia with a single administration. However, the therapeutic effect of hemophilia A gene therapy is not long-lasting and the amount of vector required is high. One possible solution involves the modification of FVIII protein; the functional engineering of FVIII may lead to stable gene therapy at low doses. Several engineered FVIII variants that use amino acid substitutions from pigs and other animal species have been reported^21,27^. In the present study, our engineered FVIII improved FVIII secretion, increased FVIII cofactor activity, and allowed for treatment at much lower doses than previous hemophilia A gene therapy.

Gene therapy for hemophilia A using AAV vectors is advancing in clinical trials with various serotypes; however, these trials require higher doses of vectors than therapies for hemophilia B. Valoctocogene roxaparvovec is a gene therapy product that uses the AAV5 serotype; it is approved by both the US Food and Drug Administration and the European Medicines Agency. In a phase 1/2 study of valoctocogene roxaparvovec at the same vector dose (6 × 10^12^ vg/kg) at which FVIII exceeded physiological concentrations in our macaque study, plasma FVIII was less than 1%^1^. An AAV8 vector carrying the modified FVIII, which included a 17-amino-acid peptide with six *N*-linked glycosylation motifs from the human FVIII B-domain (FVIII-V3), achieved three times higher FVIII expression in animal models compared with FVIIISQ^4^. Moreover, in a phase 1/2 trial (NCT03001830) using FVIII-V3, two patients received a dosage of 2 × 10^12^ vg/kg, which resulted in FVIII activity levels of 6 and 69 IU/dL^39^; these levels are lower than those from our engineered FVIII in macaques. Gene therapy for hemophilia A using our engineered FVIII may therefore achieve sufficient therapeutic effects with vector doses comparable with or lower than those used in hemophilia B gene therapy, even when wild-type AAV serotypes are used.

Because our engineered FVIII potentially allows for a reduction in the AAV vector dose, both vector-related hepatotoxicity and immune-related adverse reactions may be reduced. Although AAV vectors have generally been considered to have low immunogenicity, recent findings from clinical trials targeting various diseases have highlighted that immune-related adverse reactions are an important concern. These reactions involve both innate immunity and T-cell-mediated acquired immunity. Furthermore, thrombotic microangiopathy and liver damage resulting from immune reactions to AAV increase with higher vector doses^13^. Additionally, in hemophilia gene therapy, immune modulation with corticosteroids is often required when relatively high doses of AAV vectors are used^40^. Our engineered FVIII variant may thus facilitate safer hemophilia A gene therapy by minimizing immune reactions to AAV and producing effective treatment at lower vector doses, even when conventional serotypes are used. Engineered AAV serotypes with higher liver tropism are currently being used in hemophilia gene therapy^41,42^; combining these modified serotypes with our FVIII variant may further reduce the vector dose. Notably, achieving effective therapy at lower vector doses will also be advantageous for reducing the cost of vector production.

Our engineered FVIII variant enhanced FVIII secretion in hepatocytes and reduced intracellular FVIII accumulation. FVIII overexpression triggers protein aggregation in the ER and activates UPR^8,43^. Addressing UPR induction during FVIII expression is thus a challenge that must be resolved to achieve long-term efficacy of hemophilia A gene therapy. Binding immunoglobulin protein/Glucose-regulated protein (BiP/GRP78), which prevents protein aggregation in the ER, binds to unfolded/misfolded proteins and releases them in an ATP-dependent manner^44^. Consequently, FVIII secretion requires substantial amounts of ATP^43^. High FVIII expression can deplete cellular ATP and further promote the formation of high-molecular-weight aggregates^45^. The aggregation motifs in the A1 domain are crucial for intracellular FVIII accumulation^46^. Indeed, amino acid substitutions in the A1 domain, such as in variant X5, enhance secretion^21^. In addition, the transport of FVIII from the ER to the Golgi requires the binding of *N*-linked glycans of FVIII with the cargo receptor complex lectin, mannose binding 1/multiple coagulation factor deficiency 2 (LMAN1/MCFD2)^47^. FVIII-V3, which retains six *N*-glycosylation sites, was developed to target this mechanism by adding glycan modifications to enhance secretion^8^. Our engineered FVIII incorporates a new *N*-linked glycan (K213N) in the A1 domain, which may play an important role in promoting secretion. However, modification of the heavy chain alone is likely insufficient for efficient secretion because proper heterodimer formation with the light chain is crucial for FVIII secretion^48^. In our engineered variant, the reduced hepatocytic accumulation of the light chain suggests that the introduction of multiple amino acid substitutions in the light chain enhances its association with the heavy chain, thereby improving secretion.

It should be noted that the mechanisms of FVIII secretion may vary depending on the cell type. In gene therapy for hemophilia, hepatocytes are predominantly targeted because of high AAV receptor expression and gene transfer efficiency. However, FVIII is produced physiologically by sinusoidal endothelial cells. FVIII then associates with von Willebrand factor (VWF) secreted from endothelial cells and is stabilized in the blood. Although it remains unclear whether VWF binds FVIII intracellularly and secretes it under physiological conditions, the simultaneous gene transfer of FVIII with VWF alters the intracellular transport of FVIII, leading to its storage and co-localization with VWF in granules^49^. Additionally, the gene transfer of FVIII into endothelial cells results in the storage of FVIII and VWF in Weibel–Palade bodies^49^. The intracellular interactions between FVIII and VWF, and their secretion from Weibel–Palade bodies, may represent a regulated physiological pathway for FVIII secretion. Therefore, for gene therapy targeting hemophilia A, approaches such as gene transfer into sinusoidal endothelial cells—or genome editing therapies targeting these cells—are potential strategies for achieving efficient FVIII secretion.

Our engineered FVIII variant exhibited high specific coagulation factor activity. This increased specific activity was likely caused by its high affinity for FIXa, as indicated by our biochemical and structural analyses. This may then facilitate the early generation of FXa. Furthermore, our variant showed a significant discrepancy between the one-stage clotting assay and the chromogenic assay; the one-stage assay yielded substantially higher activity. The chromogenic substrate assay measures FVIIIa activity after a specified period, whereas the one-stage clotting assay evaluates total thrombin generation from the initial coagulation reaction^50^. Notably, hemophilia patients with point mutations sometimes exhibit discrepancies between the two assays. Elevated FVIII activity in the one-stage assay is thought to be associated with point mutations in the hydrophobic core located at the A1–A2–A3 domain interface^51^. This observation is consistent with the increased A2 domain dissociation in our engineered variant, which is associated with reduced non-covalent interactions at the A1–A2–A3 domain interface. Together, these findings suggest that the discrepancy in FVIII activity between the two assays in our variant was likely caused by the early, strong generation of FXa by FIXa, as well as the rapid dissociation of the A2 domain. Although sustained FVIII activation might increase the risk of thrombosis, our engineered variant may produce an enhanced hemostatic response while potentially reducing the risk of thrombosis, even at high expression levels.

Our study has several limitations. First, there is the issue of immunogenicity. In our experiments with macaques, antibodies against human FVIII were generated. In clinical hemophilia gene therapy, eligible patients can develop immune tolerance through the repeated administration of a product, which minimizes the risk of inhibitor formation. Although our engineered variant did not show any new epitopes with high antigenicity in the *in silico* analyses, the amino acid substitutions from the human sequence warrant careful observation to avoid inhibitor formation. Second, we have not yet elucidated which of the introduced amino acid substitutions contributes to the increased activity, improved secretion, and/or reduced UPR; further molecular structural evaluations are necessary. Finally, the emergence of inhibitors against human FVIII prevented long-term observations in our macaque trials. The persistence of FVIII expression thus needs to be carefully observed in human clinical trials.

We have successfully developed an engineered FVIII that may lead to effective gene therapy against hemophilia A. Currently, novel therapies such as extended half-life products, bispecific antibodies, and anti-tissue factor pathway inhibitor antibodies have substantially improved patient outcomes and quality of life. Unlike existing treatments, gene therapy offers the potential for patients to avoid being constantly aware of their hemophilia because only a single treatment is required. The World Federation of Hemophilia reported that 270,000 individuals have hemophilia worldwide, of whom 85% are not receiving adequate treatment^52^. Gene therapy is thus also a promising treatment option in regions with limited medical access. Moreover, the advancement of gene therapy is expected to benefit not only patients but also female carriers, affecting life events such as marriage and childbirth. Achieving effective and safe gene therapy therefore holds important potential to improve the quality of life of hemophilia patients and their families. Moving forward, we aim to advance into human clinical trials and contribute to the treatment of hemophilia patients and their families.

## Online Methods

### Cell culture

HEK293 cells (JRCB Cell Bank, Osaka, Japan), and AAVpro 293T cells (Takara Bio, Shiga, Japan) were cultured in Dulbecco’s modified Eagle Medium (Sigma Aldrich, Saint Louis, MO) supplemented with 10% fetal bovine serum (Thermo Fisher Scientific, Waltham, MA) and GlutaMAX^TM^ Supplement (Thermo Fisher Scientific). Huh-7 cells (Institute of Development, Aging and Cancer, Tohoku University, Miyagi, Japan) were cultured in RPMI-1640 Medium (Sigma Aldrich) supplemented with 10% fetal bovine serum and GlutaMAX Supplement. To select the transfected cell clones, G418 (Nacalai Tesque Inc., Kyoto, Japan) was added to the culture medium; the culture was continued for selection. To isolate and purify FVIIISQ proteins, the stable cell lines were acclimated to serum-free HE100 medium (GMEP Cell Technologies, Fukuoka, Japan) supplemented with GlutaMAX Supplement, and the culture supernatant was collected to purify FVIIISQ proteins. For Huh-7 cells, the culture supernatant used for immunoblotting was serum-free ISE-RPMI medium containing 5 mM HEPES, 30 nM Na_2_SeO_3_, 3 nM (NH_4_)_6_Mo_7_O_2_·4H_2_O, 10 μM FeSO_4_·7H_2_O, 0.3 nM MnCl_2_·4H_2_O, 10 nM NH_4_Vo_3_, 3 nM linoleic acid, 3 nM oleic acid, 30 μM ethanolamine, and 24 mM NaHCO_3_ in RPMI-1640, as previously reported^53^.

### cDNA

The cDNA encoding human B-domain deleted FVIIISQ (hFVIIISQ) was codon-optimized and synthesized using GeneArt (Thermo Fisher Scientific) with the amino acid sequences (NP_000123.1) as previously described^38^. The cDNAs of canine FVIIISQ (cFVIIISQ) (NP_001003212.1), porcine FVIIISQ (pFVIIISQ) (NP_999332.2), bovine FVIIISQ (bFVIIISQ) (XP_024843485.1), and ovine FVIIISQ (oFVIIISQ) (NP_001166021.1) were codon-optimized and synthesized using GeneArt. The B-domain was replaced into the common linker sequence SFSQNPPVLKRHQR. Because the synthesized cDNA of cFVIIISQ did not express functional FVIII in the supernatant, we induced K461N, L574F, and D584N based on the UniProt database (O18806). FVIIISQ(Ver.1) was the engineered hFVIIISQ in which we replaced 72 amino acids that were common to cFVIIISQ and pFVIIISQ but different in hFVIIISQ. In FVIIISQ(Ver.2), we replaced 58 amino acids that were common to cFVIIISQ, pFVIIISQ, bFVIIISQ, and oFVIIISQ but different in hFVIIISQ. FVIIISQ(Ver.3) was created based on FVIIISQ(Ver.1), but the 21 amino acids in FVIIISQ(Ver.1) that had unchanged physiochemical properties (e.g., hydrophilic and negatively charged, D and E; hydrophilic, S and T; hydrophilic and positively charged, K, H, and R; hydrophobic and aliphatic, I, L, and V) were restored to the original hFVIIISQ sequences (51 amino acid substitutions remained). In FVIIISQ(Ver.4), we replaced 36 amino acids that were common to FVIIISQ(Ver.2) and FVIIISQ(Ver.3). ET3, a chimeric FVIII of hFVIIISQ and pFVIIISQ, was produced by replacing the A1–a1 and a3–A3 domains of pFVIIISQ with hFVIIISQ^54^. The A1 domain DNA fragment of FVIIISQ-X5^21^ and the A3–C1 domain DNA fragment of FVIIISQ-JF12^27^ were synthesized by GenScript Biotech Corp (Piscataway, NJ) and Eurofins Genomics (Luxembourg, Luxembourg), respectively. The fragment was inserted into hFVIIISQ using an In-Fusion HD Cloning Kit (Takara Bio). The FVIIISQ(Ver. 4) without CpG sequences was designed and synthesized by GeneScript Japan, Azenta (Tokyo, Japan), and Fasmac (Kanagawa, Japan). The mTTR proximal promoter containing a distal enhancer^55^ was synthesized by Eurofins Genomics.

### Plasmid transfection

To screen for FVIII expression in Huh-7 cells, FVIIISQ cDNAs were inserted into a plasmid containing the HCRhAAT chimeric promoter (an enhancer element of the hepatic control region of the Apo E/C1 gene and the human anti-trypsin promoter) and SV40 polyadenylation signals. For stably FVIIISQ-expressing transfectants, the FVIIISQ cDNAs were incorporated into a pcDNA3 plasmid (Thermo Fisher Scientific). The plasmid was transfected into the cells using Lipofectamine 3000 Reagent (Thermo Fisher Scientific) according to the manufacturer’s recommendations.

### Measurements of FVIII:C, FVIII:Ag, and FVIII inhibitor

FVIII:C was measured using the one-stage clotting assay based on activated partial thromboplastin time (Sysmex, Hyogo, Japan) and a chromogenic assay based on a chromogenic substrate (Revohem FVIII Chromogenic, a blood coagulation FVIII measurement kit; Sysmex) with an automated coagulation analyzer (CS-1600; Sysmex). Human FVIII:Ag in the supernatant and mouse plasma was measured using Asserachrom VIII:Ag (Stago, Asnières sur Seine, France) according to the manufacturer’s instructions.

To detect human FVIII:Ag in monkey plasma, we developed a polyclonal anti-hFVIII antibody from a monkey immunized with recombinant hFVIII (Turoctocog Alpha, Novo Nordisk A/S, Bagsvaerd, Denmark). The serum was obtained from the monkey, whose serum inhibited human FVIII:C but not monkey FVIII:C. Immunoglobulin G (IgG) was then isolated from the serum using a Melon Gel IgG Purification Kit (Thermo Fisher Scientific). The antibody that specifically bound to hFVIII was isolated using column chromatography (AminoLink Plus Immobilization Kit, Thermo Fisher Scientific) conjugated with recombinant hFVIII. The isolated antibody against hFVIII was coated in 96-well plates (0.5 μg/mL) overnight at 4°C. The samples were blocked with 5% casein in phosphate-buffered saline (PBS) for 1 hour at room temperature, and were then incubated in PBS containing 1% casein and 0.1% Triton X-100 for 1 hour at 37°C. After washing with PBS containing 0.1% Triton X-100, the bound FVIII antigen was reacted with 1 μg/mL of anti-hFVIII antibody (GMA8023, Green Mountain Antibodies, Burlington, VT). Antibody binding was detected using anti-mouse IgG conjugated with horseradish peroxidase (Seracare, Milford, MA) and a TMB Substrate Kit (Thermo Fisher Scientific).

The presence of FVIII-neutralizing antibodies (i.e., inhibitor) was identified using the Bethesda method^56^. Normal pooled plasma (as a source of FVIII) was incubated with a heat-inactivated plasma sample for 2 hours. The residual FVIII:C was then measured using the one-stage clotting assay. One Bethesda unit/mL was defined as the amount of inhibitor that inhibited 50% of the FVIII:C.

### Evaluation of UPR

The XBP-luciferase gene, which expresses luciferase during IRE1-induced UPR^57^, was constructed by conjugating the luciferase gene with *XBP1* (GeneScript Japan). The conjugated gene (*XBP*-luciferase) was then cloned into pBApo-EF1α-Neo (Takara Bio). Next, the linearized plasmid was transfected into Huh-7 cells using Lipofectamine 3000. After selection using G418 treatment, we isolated a single clone that had low baseline luciferase activity but high luciferase activity with the addition of tunicamycin (FUJIFILM Wako Pure Chemical Corporation, Osaka, Japan). The Huh-7 cells stably expressing XBP-luciferase were seeded into 24-well plates at 1.5 × 10^5^ cells/well before being transfected with the plasmid using Lipofectamine 3000. After 48 hours of incubation, the cells were lysed using 1× Passive Lysis buffer (Promega, Madison, WI). Luciferase activity was then measured using a luminometer (Centro LB960, Berthold Technologies, Bad Wildbad, Germany), as described previously^58^.

### Immunoblotting of FVIII

Huh-7 cells were transfected with the plasmid, and the culture medium was replaced with serum-free medium the day after transfection. The medium was collected 24 hours later and centrifuged at 2500 × *g* for 5 min. The cells were then lysed in PBS containing 0.5% Triton X-100, and the lysate was centrifuged at 20,000 × *g* for 10 min. The supernatant was recovered as the cell lysate. The supernatants and cell lysates were resolved by sodium dodecyl sulfate polyacrylamide gel electrophoresis and transferred to a polyvinylidene fluoride membrane. The membranes were blocked with 5% (weight/volume) skim milk powder in 20 mM Tris (pH 7.5), 150 mM NaCl, and 0.05% Tween 20 (TBS-T buffer) for 1 hour. After extensive washing with TBS-T, the membranes were incubated overnight at 4°C with an anti-hFVIII polyclonal antibody (#PAHFVIII-S, 1/20,000 dilution, Haematologic Technologies Inc., Essex Junction, VT) in TBS-T containing 5% (weight/volume) bovine serum albumin. Antibody binding was detected using horseradish peroxidase (HRP)-conjugated anti-sheep IgG (Bethyl Laboratories, Inc., Montgomery, TX) and visualized with Immobilon Western Chemiluminescent HRP Substrate (Millipore, Burlington, MA) and an ImageQuant LAS4000 digital imaging system (GE Healthcare, Buckinghamshire, UK). The bands in the captured images were quantified using ImageJ.

### AAV vector production

A DNA fragment consisting of a chimeric HCRhAAT promoter or mTTR promoter, FVIIISQ cDNAs, and the SV40 polyadenylation signal was introduced into the pAAV plasmid between AAV2-derived inverted terminal repeats. The AAV genes were packaged using the triple plasmid transfection of AAVpro293T cells to generate the AAV vector (helper-free system), as described previously^59^. The pHelper plasmid (Takara Bio), the pRC8 plasmid expressing Rep and serotype 8 capsid (AAV8) or the pRC5 plasmid (Takara Bio) expressing Rep and serotype 5 capsid (AAV5), and the pAAV plasmid containing a gene of interest were simultaneously transfected; the AAV vectors were purified from the transfected cells 72 hours after transduction, as described previously^60^. The titration of recombinant AAV vectors was performed using quantitative PCR (qPCR) targeting the SV40 polyA sequence^59^. The primer and probe sequences are described in Supplementary Table 6.

### Animal experiments

All experimental animal procedures were approved by The Institutional Animal Care and Concern Committee of Jichi Medical University (permission numbers: 20051-09, 23062-01, and 23067-01) and Tsukuba Primate Research Center (DSR03-15). Animal care was conducted in accordance with the committee’s guidelines and the ARRIVE (Animal Research: Reporting of In Vivo Experiments) guidelines^61^. The FVIII-deficient mice (B6;129S4-*F8^tm1Kaz^*/J) were kindly provided by Dr. H.H. Kazazian Jr. (University of Pennsylvania, Philadelphia, PA). Animals were maintained in isolators in the specific pathogen-free facility of Jichi Medical University at 23°C ± 3°C with 12:12-hour light/dark cycles. To obtain plasma samples, mice were anesthetized with isoflurane (1%–3%) and blood samples were drawn from the jugular vein using a 29G micro-syringe (TERUMO, Tokyo, Japan) containing 1/10 (volume/volume) sodium citrate (Harasawa Pharmaceutical, Tokyo, Japan). Plasma was isolated by centrifugation before being frozen and stored at −80°C until its analysis. The AAV vector was administered intravenously through the jugular vein (100−150 µL).

The intravenous injection of the AAV vector into macaques (*Macaca fascicularis*) was performed at Tsukuba Primate Research Center, National Institutes of Biomedical Innovation, Health and Nutrition (Ibaraki, Japan). We selected seven male macaques without anti-AAV neutralizing antibodies against AAV8 or AAV5 (Supplementary Table 7). The AAV vector was intravenously administered for 5 min via a saphenous vein. To reduce vector immunogenicity, prednisolone (1 mg/kg/day) and tacrolimus hydrate (0.05 mg/kg/day) were intramuscularly administered for 56 and 28 days, respectively. Prednisolone was then gradually reduced (0.5 mg/kg/day [days 57–63], 0.3 mg/kg/day [days 64–70], 0.2 mg/kg/day [days 71–77], and 0.1 mg/kg/day [days 78–84]).

### qPCR

Genomic DNA and RNA were extracted using a DNeasy Blood & Tissue Kit (QIAGEN, Venlo, Netherlands) and an RNeasy Mini Kit (QIAGEN), respectively. The RNA samples were reverse-transcribed using a PrimeScript RT Reagent Kit (Takara Bio). qPCR was performed using THUNDERBIRD™ Probe qPCR Mix (TOYOBO, Osaka, Japan) or THUNDERBIRD SYBR qPCR Mix (TOYOBO) with a QuantStudio 12K Flex (Thermo Fisher Scientific). Quantification of the AAV genome into genomic DNA was measured by targeting the SV40 polyA sequence. mRNA expression levels were expressed as copy numbers in 100 ng RNA. Copy numbers were calculated using a known copy number of the plasmid harboring the target sequence. Where indicated, digital PCR was used to quantify the AAV vector genome in organs (QuantStudio Absolute Q Digital PCR System, Thermo Fisher Scientific). The primer and probe sequences are described in Supplementary Table 6.

### Protein preparation and biochemical analysis of FVIII

FVIIISQ(Ver.4) and wild-type FVIIISQ proteins were purified from the supernatants of stably expressing HEK293 cells. The culture supernatant from stably expressing cells cultured in serum-free medium (HE100 medium) was filtered through a 0.22 µm filter after centrifugation. The FVIII proteins in the culture supernatant were then purified using affinity chromatography with VIIIselect (Cytiva, Marlborough, MA) and size exclusion chromatography. Purified FVIII concentrations were measured using enzyme-linked immunosorbent assay with two anti-FVIII monoclonal antibodies (anti-FVIII C2; ESH8 and anti-FVIII A2; R8B12)^62^. Concentrations were verified by measuring A_280_ absorbance^63^. To calculate the *K*_m_ and *K*_cat_ for the FX–FVIIIa association and the apparent *K*d and *V*_max_ for the FIXa–FVIIIa association, the rate of FX conversion to FXa was monitored using a purified system, as previously reported^64^. The thrombin generation assay, triggered by recombinant tissue factor, was performed as previously described^64^.

### Single-particle cryo-electron microscopy

Purified FVIIISQ(Ver.4) was concentrated (A_280 nm_ = 1.5) and applied to Au 300-mesh R1.2/1.3 grids (Quantifoil, Großlöbichau, Germany) that were glow-discharged after adding 3 μL amylamine in a Vitrobot Mark IV (Thermo Fisher Scientific) at 4°C, with a waiting time of 10 s and a blotting time of 5 s under 100% humidity conditions. The grids were then plunge-frozen in liquid ethane and cooled to the temperature of liquid nitrogen. Micrographs for all datasets were collected using a Titan Krios G3i microscope (Thermo Fisher Scientific) running at 300 kV and equipped with a GIF Quantum LS Energy Filter (Gatan, Pleasanton, CA) and a K3 Summit direct electron detector (Gatan) in electron counting mode (The University of Tokyo, Japan). Datasets were collected with a total dose of approximately 50 electrons per Å^2^ per 48 frames, using the standard mode. The dose-fractionated movies were processed using cryoSPARC v3.3.2. For data processing details, see Supplementary Fig. 5.

### Model building and validation

The model was built using the predicted model of the FVIIISQ(Ver.4) protein created by AlphaFold 2^65^ as the reference, followed by manual model building with Coot^66^ against the density map. The model was refined using Phenix 1.21.2.5419^67^. Residues 1–18, 40–61, 128–130, 353–395, 732–818, and 840–849 were not included in the final model because of their poor resolutions in the density map. All molecular graphics in Fig. 4 were prepared using UCSF ChimeraX^68^.

### *In silico* structural analysis

We used AlphaFold 3^69^ to generate complete structural models of FVIII by complementing disordered regions in the cryo-electron microscopy structures. We selected high-ranking models and confirmed structural alignment using the TM-align algorithm^70^. We used AlphaFold-Multimer to predict the potential binding interactions between FVIII and FIXa^69^. To quantify the binding affinity between FVIII and the protease domain of FIXa, we used HADDOCK 2.4^71^. The results were expressed as HADDOCK scores, electrostatic energy, Van der Waals energy, and buried surface area to determine the strength and stability of the FVIII–FIXa complexes. Protein structures were visualized and analyzed using PyMOL molecular visualization software (version 2.6, Schrödinger, Inc., New York, NY). Electrostatic potential calculations of the molecular surface were performed using the PyMOL APBS plugin with the default settings, which implements the Adaptive Poisson–Boltzmann Solver^72^.

### Post-translational modification

FVIIISQ(Ver.4) protein was denatured, reduced, and alkylated using 6 M Gdn-HCl, 5 mM tris(2-carboxyethyl) phosphine, and 50 mM iodoacetamide in the presence of 50 mM ammonium bicarbonate at 37°C in the dark for 1 hour. Desalting was performed using Zeba Spin Desalting Columns (Thermo Fisher Scientific) according to the manufacturer’s instructions. The sample was digested with trypsin at a 1:20 enzyme/substrate ratio at 37°C overnight. Digestion was then halted by adding 10% formic acid, and the sample was vacuum dried, dissolved in 0.1% formic acid in water, and analyzed using liquid chromatography–tandem mass spectrometry (LC-MS/MS). Two different fragmentation methods were used in the LC-MS/MS analysis: higher-energy collisional dissociation (HCD) and electron-transfer/higher-energy collisional dissociation (EThcD).

The LC-MS/MS analysis using HCD fragmentation was performed using Ultimate 3000 (Thermo Fisher Scientific) LC pumps coupled to a Q Exactive HF-X mass spectrometer (Thermo Fisher Scientific), with a mobile phase A of 0.1% formic acid in water and a mobile phase B of 0.1% formic acid in acetonitrile. Peptides were trapped to a PepMap300 C18 column (1.0 × 15 mm, 5 µm particle size; Thermo Fisher Scientific), and separated using an ACQUITY UPLC BEH C18 column (1.0 × 100 mm, 1.7 µm particle size; Waters Corporation, Milford, MA). The LC conditions were as follows: a gradient of 0%–7% B for 4 minutes, 7%–10% B for 6 minutes, 10%–25% B for 30 minutes, 25%–35% B for 15 minutes, and 35%–95% B for 5 minutes. The flow rate was set at 50 µL/min. A full MS scan was performed at a spray voltage of 3.8 kV, a capillary temperature of 300°C, a resolution of 60,000, a mass range (*m*/*z*) of 300–2000, and a funnel radio frequency level of 40. A subsequent data-dependent MS/MS scan was performed at a higher energy collisional dissociation of 27% and a resolution of 15,000.

The LC-MS/MS analysis using EThcD fragmentation was performed using Vauquish Neo (Thermo Fisher Scientific) LC pumps coupled to a Orbitrap Eclipse mass spectrometer (Thermo Fisher Scientific), with a mobile phase A of 0.1% formic acid in water and a mobile phase B of 0.1% formic acid in acetonitrile. Peptides were separated using an ACQUITY UPLC CSH C18 column (1.0 × 150 mm, 1.7 µm particle size; Waters Corporation, Milford, MA). The LC conditions were as follows: a gradient of 5%– 40% B for 50 minutes and 40%–95% B for 2 minutes. The flow rate was set at 50 µL/min. A full MS scan was performed at a spray voltage of 3.5 kV, a capillary temperature of 300°C, a resolution of 60,000, a mass range (*m*/*z*) of 350–1800, and a funnel radio frequency lens value of 30. A subsequent data-dependent MS/MS scan was performed at a higher energy collisional dissociation of 30% and a resolution of 30,000. EThcD mass trigger was set to the typical glycan ions 138.0545, 204.0867, and 366.1396 with mass tolerance of 15 ppm. ETD reagent target was set at 2.0e5, and ETD reaction time was set at 60 ms, 40 ms, and 20 ms for the charge states 2, 3, and 4 to 8, respectively.

The data analysis was performed using Proteome Discoverer (version 2.4, Thermo Fisher Scientific).

### *In silico* analysis

We used NetMHCIIpan^33^ to estimate the binding affinities of all possible engineered FVIIISQ(Ver.4) sequences to each of the 46 selected HLA subtypes (i.e., the *DRB1*, *DRB3*, *DRB4*, *DRB5*, *DP*, and *DQ* groups). For this purpose, we used all fragments sized between 15 and 20 of the wild-type FVIIISQ and engineered FVIIISQ(Ver.4) sequences. From the program output, we selected only the epitopes that: (i) did not exist in the wild-type FVIIISQ; and (ii) had a predicted binding affinity < 500 nM. In NetMHCIIpan, epitopes have a core region and surrounding residues. For this reason, some epitopes had multiple predicted binding affinities; in such cases, we considered the strongest binding affinity for each epitope. Scripts were written in Python (version 3.8), and additional analyses were conducted using the R statistical package (version 4.4.1).

### Statistical analysis

Statistical analyses were performed using GraphPad Prism^®^ (GraphPad Software, San Diego, CA). All data are presented as the standard error of the mean (SEM) or standard deviation (SD). The statistical methods in each experiment are described in the figure legends.

## Acknowledgments

This work was supported by AMED (21fk0410037, 24fk0410061, 24bk0304007, 23ae0201007, and 24bm1323001), Japan, the Research Grant program by the Japanese Society of Thrombosis and Hemostasis, and the SENSHIN Medical Research Foundation. Optima XE-90 was subsidized by JKA through its promotion funds from KEIRIN RACE. We thank Yaeko Suto, Mika Kishimoto, Tamaki Aoki, Sachiyo Kamimura, Mai Hayashi, Yuiko Ogihara, Nagako Sekiya, Noguchi Tomoko, Hiromi Ozaki, and Hiroko Hayakawa of Jichi Medical University for their technical assistance. We thank Yoshiki Nagashima and Daisuke Higo of Thermo Fisher Scientific for EThcD analysis of PTM. We also thank Bronwen Gardner, PhD, from Edanz (https://jp.edanz.com/ac) for editing a draft of this manuscript.

## Author contributions

Y.Kash. and T.O. conceived the study. Y.Kash. and T.O. wrote the manuscript. Y.N. and K.N. performed the biochemical analysis of FVIII. Y.F., Y.Y., and S.U. performed the translational modification analysis. K.H. and O.N. performed the structural analysis by cryo-electron microscopy. N.B. performed the structural-based prediction analysis. T.L performed the *in silico* analysis. M.H. analyzed the data and revised the manuscript. Y.Kata. performed the non-human primate experiments. All authors provided feedback on the manuscript.

## Corresponding author

Correspondence to Yuji Kashiwakura and Tsukasa Ohmori. Lead contact: Tsukasa Ohmori

## References

1. Rangarajan, S. et al. AAV5–Factor VIII Gene Transfer in Severe Hemophilia A. New England Journal of Medicine 377, 2519–2530 (2017).

2. George, L. A. et al. Multiyear Factor VIII Expression after AAV Gene Transfer for Hemophilia A. New England Journal of Medicine 385, 1961–1973 (2021).

3. Pasi, K. J. et al. Multiyear Follow-up of AAV5-hFVIII-SQ Gene Therapy for Hemophilia A. N Engl J Med 382, 29–40 (2020).

4. McIntosh, J. et al. Therapeutic levels of FVIII following a single peripheral vein administration of rAAV vector encoding a novel human factor VIII variant. Blood 121, 3335–3344 (2013).

5. Ozelo, M. C. et al. Valoctocogene Roxaparvovec Gene Therapy for Hemophilia A. N Engl J Med 386, 1013–1025 (2022).

6. Samelson-Jones, B. J. & George, L. A. Adeno-Associated Virus Gene Therapy for Hemophilia. Annu. Rev. Med. 74, 231–247 (2023).

7. Hayakawa, M. et al. Characterization and visualization of murine coagulation factor VIII-producing cells in vivo. Sci Rep 11, 14824 (2021).

8. Malhotra, J. D. et al. Antioxidants reduce endoplasmic reticulum stress and improve protein secretion. Proc. Natl. Acad. Sci. U.S.A. 105, 18525–18530 (2008).

9. Brown, H. C., Gangadharan, B. & Doering, C. B. Enhanced Biosynthesis of Coagulation Factor VIII through Diminished Engagement of the Unfolded Protein Response. Journal of Biological Chemistry 286, 24451–24457 (2011).

10. Amen, O. M., Sarker, S. D., Ghildyal, R. & Arya, A. Endoplasmic Reticulum Stress Activates Unfolded Protein Response Signaling and Mediates Inflammation, Obesity, and Cardiac Dysfunction: Therapeutic and Molecular Approach. Front Pharmacol 10, 977 (2019).

11. Fong, S. et al. Interindividual variability in transgene mRNA and protein production following adeno-associated virus gene therapy for hemophilia A. Nat Med 28, 789–797 (2022).

12. Lenting, P. J., Denis, C. V. & Christophe, O. D. Emicizumab, a bispecific antibody recognizing coagulation factors IX and X: how does it actually compare to factor VIII? Blood 130, 2463–2468 (2017).

13. Mullard, A. Gene therapy community grapples with toxicity issues, as pipeline matures. Nat Rev Drug Discov 20, 804–805 (2021).

14. Wang, J.-H., Gessler, D. J., Zhan, W., Gallagher, T. L. & Gao, G. Adeno-associated virus as a delivery vector for gene therapy of human diseases. Sig Transduct Target Ther 9, 78 (2024).

15. George, L. A. et al. Hemophilia B Gene Therapy with a High-Specific-Activity Factor IX Variant. N Engl J Med 377, 2215–2227 (2017).

16. Simioni, P. et al. X-Linked Thrombophilia with a Mutant Factor IX (Factor IX Padua). New England Journal of Medicine 361, 1671–1675 (2009).

17. Pipe, S. W. & Kaufman, R. J. Characterization of a genetically engineered inactivation-resistant coagulation factor VIIIa. Proc. Natl. Acad. Sci. U.S.A. 94, 11851–11856 (1997).

18. Wilhelm, A. R. et al. Activated protein C has a regulatory role in factor VIII function. Blood 137, 2532–2543 (2021).

19. Siner, J. I., et al. Circumventing furin enhances factor VIII biological activity and ameliorates bleeding phenotypes in hemophilia models. JCI Insight 1, (2016).

20. Nguyen, G. N. et al. Altered cleavage of human factor VIII at the B-domain and acidic region 3 interface enhances expression after gene therapy in hemophilia A mice. Journal of Thrombosis and Haemostasis 21, 2101–2113 (2023).

21. Cao, W. et al. Minimal Essential Human Factor VIII Alterations Enhance Secretion and Gene Therapy Efficiency. Molecular Therapy - Methods & Clinical Development 19, 486–495 (2020).

22. Doering, C. B., Healey, J. F., Parker, E. T., Barrow, R. T. & Lollar, P. High Level Expression of Recombinant Porcine Coagulation Factor VIII. Journal of Biological Chemistry 277, 38345–38349 (2002).

23. Doering, C. et al. Expression and characterization of recombinant murine factor VIII. Thromb Haemost 88, 450–458 (2002).

24. Zakas, P. M. et al. Development and Characterization of Recombinant Ovine Coagulation Factor VIII. PLoS ONE 7, e49481 (2012).

25. Sabatino, D. E. et al. Recombinant canine B-domain–deleted FVIII exhibits high specific activity and is safe in the canine hemophilia A model. Blood 114, 4562–4565 (2009).

26. Brown, H. C. et al. Bioengineered coagulation factor VIII enables long-term correction of murine hemophilia A following liver-directed adeno-associated viral vector delivery. Molecular Therapy - Methods & Clinical Development 1, 14036 (2014).

27. Firrman, J. et al. Identification of Key Coagulation Activity Determining Elements in Canine Factor VIII. Molecular Therapy - Methods & Clinical Development 17, 328–336 (2020).

28. Lee, K. et al. IRE1-mediated unconventional mRNA splicing and S2P-mediated ATF6 cleavage merge to regulate XBP1 in signaling the unfolded protein response. Genes Dev. 16, 452–466 (2002).

29. Mellquist, J. L., Kasturi, L., Spitalnik, S. L. & Shakin-Eshleman, S. H. The amino acid following an asn-X-Ser/Thr sequon is an important determinant of N-linked core glycosylation efficiency. Biochemistry 37, 6833–6837 (1998).

30. Fay, P. J. Activation of factor VIII and mechanisms of cofactor action. Blood Reviews 18, 1–15 (2004).

31. Valley, C. C. et al. The methionine-aromatic motif plays a unique role in stabilizing protein structure. J Biol Chem 287, 34979–34991 (2012).

32. Childers, K. C., Peters, S. C. & Spiegel Jr, P. C. Structural insights into blood coagulation factor VIII: Procoagulant complexes, membrane binding, and antibody inhibition. Journal of Thrombosis and Haemostasis 20, 1957–1970 (2022).

33. Reynisson, B., Alvarez, B., Paul, S., Peters, B. & Nielsen, M. NetMHCpan-4.1 and NetMHCIIpan-4.0: improved predictions of MHC antigen presentation by concurrent motif deconvolution and integration of MS MHC eluted ligand data. Nucleic Acids Research 48, W449–W454 (2020).

34. McKinney, D. M. et al. A strategy to determine HLA class II restriction broadly covering the DR, DP, and DQ allelic variants most commonly expressed in the general population. Immunogenetics 65, 357– 370 (2013).

35. McGill, J. R., Simhadri, V. L. & Sauna, Z. E. HLA Variants and Inhibitor Development in Hemophilia A: A Retrospective Case-Controlled Study Using the ATHNdataset. Front. Med. 8, 663396 (2021).

36. Melief, C. J. M. & Kessler, J. H. Novel insights into the HLA class I immunopeptidome and T-cell immunosurveillance. Genome Med 9, 44 (2017).

37. Konkle, B. A. et al. BAX 335 hemophilia B gene therapy clinical trial results: potential impact of CpG sequences on gene expression. Blood 137, 763–774 (2021).

38. Ohmori, T. et al. Safety of intra-articular transplantation of lentivirally transduced mesenchymal stromal cells for haemophilic arthropathy in a non-human primate. Int J Hematol 108, 239–245 (2018).

39. Nathwani, A. C. et al. GO-8: Preliminary Results of a Phase I/II Dose Escalation Trial of Gene Therapy for Haemophilia a Using a Novel Human Factor VIII Variant. Blood 132, 489–489 (2018).

40. Nathwani, A. C. et al. Adenovirus-Associated Virus Vector–Mediated Gene Transfer in Hemophilia B. N Engl J Med 365, 2357–2365 (2011).

41. Lisowski, L. et al. Selection and evaluation of clinically relevant AAV variants in a xenograft liver model. Nature 506, 382–386 (2014).

42. Chowdary, P. et al. Phase 1–2 Trial of AAVS3 Gene Therapy in Patients with Hemophilia B. New England Journal of Medicine 387, 237–247 (2022).

43. Morris, J. A., Dorner, A. J., Edwards, C. A., Hendershot, L. M. & Kaufman, R. J. Immunoglobulin Binding Protein (BiP) Function Is Required to Protect Cells from Endoplasmic Reticulum Stress but Is Not Required for the Secretion of Selective Proteins. Journal of Biological Chemistry 272, 4327–4334 (1997).

44. Behnke, J., Mann, M. J., Scruggs, F.-L., Feige, M. J. & Hendershot, L. M. Members of the Hsp70 Family Recognize Distinct Types of Sequences to Execute ER Quality Control. Molecular Cell 63, 739–752 (2016).

45. Tagliavacca, L., Wang, Q. & Kaufman, R. J. ATP-dependent dissociation of non-disulfide-linked aggregates of coagulation factor VIII is a rate-limiting step for secretion. Biochemistry 39, 1973–1981 (2000).

46. Poothong, J. et al. Factor VIII exhibits chaperone-dependent and glucose-regulated reversible amyloid formation in the endoplasmic reticulum. Blood 135, 1899–1911 (2020).

47. Zheng, C. et al. Structural Characterization of Carbohydrate Binding by LMAN1 Protein Provides New Insight into the Endoplasmic Reticulum Export of Factors V (FV) and VIII (FVIII). Journal of Biological Chemistry 288, 20499–20509 (2013).

48. Chen, L. et al. The Enhancing Effects of the Light Chain on Heavy Chain Secretion in Split Delivery of Factor VIII Gene. Molecular Therapy 15, 1856–1862 (2007).

49. Rosenberg, J. B. et al. Intracellular trafficking of factor VIII to von Willebrand factor storage granules. J. Clin. Invest. 101, 613–624 (1998).

50. Potgieter, J. J., Damgaard, M. & Hillarp, A. One-stage vs. chromogenic assays in haemophilia A. European J of Haematology 94, 38–44 (2015).

51. Pipe, S. W., Saenko, E. L., Eickhorst, A. N., Kemball-Cook, G. & Kaufman, R. J. Hemophilia A mutations associated with 1-stage/2-stage activity discrepancy disrupt protein-protein interactions within the triplicated A domains of thrombin-activated factor VIIIa. Blood 97, 685–691 (2001).

52. *World Federation of Hemophilia Report on the Annual Grobal Survey* 2022. https://www1.wfh.org/publications/files/pdf-2399.pdf (2023).

53. Nakabayashi, H. et al. Phenotypical stability of a human hepatoma cell line, HuH-7, in long-term culture with chemically defined medium. Gan 75, 151–158 (1984).

54. Doering, C. B., Healey, J. F., Parker, E. T., Barrow, R. T. & Lollar, P. Identification of Porcine Coagulation Factor VIII Domains Responsible for High Level Expression via Enhanced Secretion. Journal of Biological Chemistry 279, 6546–6552 (2004).

55. Costa, R. H. & Grayson, D. R. Site-directed mutagenesis of hepatocyte nuclear factor (HNF) binding sites in the mouse transthyretin (TTR) promoter reveal synergistic interactions with its enhancer region. Nucl Acids Res 19, 4139–4145 (1991).

56. Yasumoto, A. et al. Overexpression of factor VII ameliorates bleeding diathesis of factor VIII-deficient mice with inhibitors. Thrombosis Research 131, 444–449 (2013).

57. Iwawaki, T., Akai, R., Yamanaka, S. & Kohno, K. Function of IRE1 alpha in the placenta is essential for placental development and embryonic viability. Proc. Natl. Acad. Sci. U.S.A. 106, 16657–16662 (2009).

58. Kashiwakura, Y. et al. Efficient gene transduction in pigs and macaques with the engineered AAV vector AAV.GT5 for hemophilia B gene therapy. Molecular Therapy - Methods & Clinical Development 30, 502–514 (2023).

59. Kashiwakura, Y. & Ohmori, T. Genome Editing of Murine Liver Hepatocytes by AAV Vector-Mediated Expression of Cas9 In Vivo. Methods Mol Biol 2637, 195–211 (2023).

60. Baatartsogt, N. et al. A sensitive and reproducible cell-based assay via secNanoLuc to detect neutralizing antibody against adeno-associated virus vector capsid. Molecular Therapy - Methods & Clinical Development 22, 162–171 (2021).

61. Percie du Sert, N., et al. The ARRIVE guidelines 2.0: Updated guidelines for reporting animal research. PLoS Biol 18, e3000410 (2020).

62. Nogami, K., Zhou, Q., Wakabayashi, H. & Fay, P. J. Thrombin-catalyzed activation of factor VIII with His substituted for Arg372 at the P1 site. Blood 105, 4362–4368 (2005).

63. Nakajima, Y., Takeyama, M., Oda, A., Shimonishi, N. & Nogami, K. Factor VIII mutated with Lys1813Ala within the factor IXa-binding region enhances intrinsic coagulation potential. Blood Advances 7, 1436–1445 (2023).

64. Nakajima, Y. et al. The combination of Asp519Val/Glu665Val and Lys1813Ala mutations in FVIII markedly increases coagulation potential. Blood Advances 8, 3929–3940 (2024).

65. Jumper, J. et al. Highly accurate protein structure prediction with AlphaFold. Nature 596, 583– 589 (2021).

66. Emsley, P. & Cowtan, K. Coot: model-building tools for molecular graphics. Acta Crystallogr D Biol Crystallogr 60, 2126–2132 (2004).

67. Afonine, P. V. et al. Real-space refinement in PHENIX for cryo-EM and crystallography. Acta Crystallogr D Struct Biol 74, 531–544 (2018).

68. Goddard, T. D. et al. UCSF ChimeraX: Meeting modern challenges in visualization and analysis. Protein Sci 27, 14–25 (2018).

69. Abramson, J. et al. Accurate structure prediction of biomolecular interactions with AlphaFold 3. Nature 630, 493–500 (2024).

70. Zhang, Y. TM-align: a protein structure alignment algorithm based on the TM-score. Nucleic Acids Research 33, 2302–2309 (2005).

71. Honorato, R. V. et al. The HADDOCK2.4 web server for integrative modeling of biomolecular complexes. Nat Protoc (2024) doi:10.1038/s41596-024-01011-0.

72. Jurrus, E. et al. Improvements to the APBS biomolecular solvation software suite. Protein Science 27, 112–128 (2018).

