## Supplementary information for "Engineered coagulation factor VIII with enhanced secretion and coagulation potential for hemophilia A gene therapy"

Yuji Kashiwakura[1,2], Yuto Nakajima[3], Kio Horinaka[4], Tiago J.S. Lopes[5], Yuma Furuta[6,7], Yuki Yamaguchi[6,7], Nemekhbayar Baatartsogt[1], Morisada Hayakawa[1,2], Yuko Katakai[8], Susumu Uchiyama[6,7], Osamu Nureki[4], Keiji Nogami[3], and Tsukasa Ohmori[1,2]

[1]Department of Biochemistry, Jichi Medical University School of Medicine, 3311-1 Yakushiji, Shimotsuke, Tochigi 329-0498, Japan

[2]Center for Gene Therapy Research, Jichi Medical University, 3311-1 Yakushiji, Shimotsuke, Tochigi 329-0498, Japan

[3]Department of Pediatrics, Nara Medical University Hospital, 840 Shijo, Kashihara, Nara 634-8522, Japan

[4]Department of Biological Sciences, Graduate School of Science, The University of Tokyo, Tokyo, Japan

[5]Nezu Life Sciences, Heidelberg, Germany

[6]Department of Biotechnology, Graduate School of Engineering, Osaka University, 2-1 Yamadaoka, Suita, Osaka 565-0871, Japan

[7]U-Medico Inc., 2-1 Yamadaoka, Suita, Osaka 565-0871, Japan

[8]The Corporation for Production and Research of Laboratory Primates, 1-16-2 Sakura, Tsukuba, Ibaraki 305-0003, Japan

Supplementary Figures: 12

Supplementary Tables: 7

### Abbreviation

FVIII, coagulation factor VIII

FVIIISQ, B-domain-deleted factor VIII

hFVIII, human FVIIISQ

cFVIII, canine FVIIISQ

pFVIII, porcine FVIIISQ

bFVIII, bovine FVIIISQ

oFVIII, ovine FVIIISQ

FVIII:C, FVIII activity

OSA, one-stage clotting assay

CSA, chromogenic substrate assay

WT, wild-type

EM, electron microscopy

3D, three-dimensional

FSC, Fourier shell correlation

AAV, adeno-associated virus

mTTRp, mouse transthyretin promoter

mTTRp_dCG, mTTRp without CpG sequences

NC, negative control

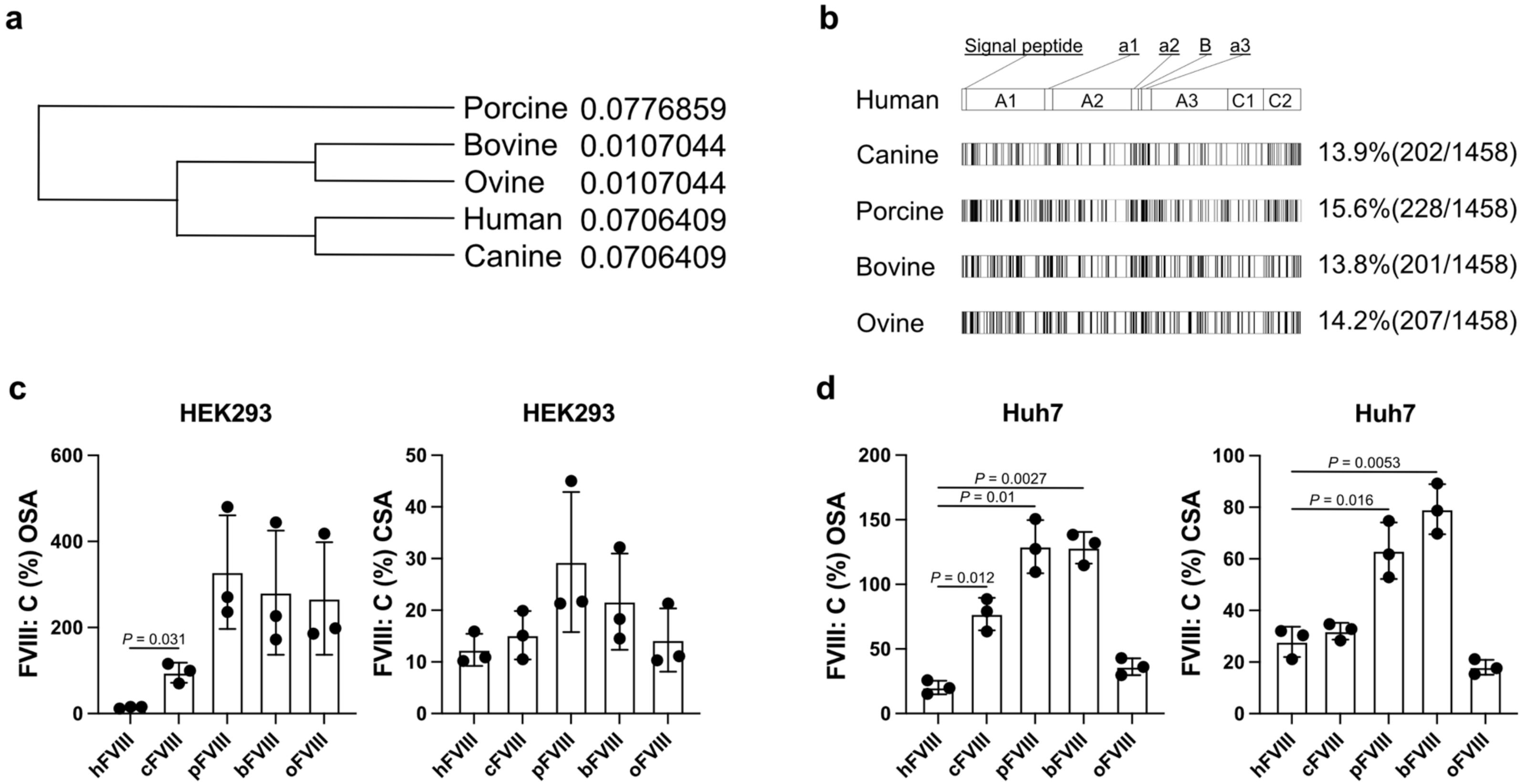

**Supplementary Fig. 1 | FVIII activity in the supernatant of cells transfected with plasmid harboring mammalian FVIIISQ.**
**(a)** Topological phylogenetic tree based on similarities in the amino acid sequences of various mammalian FVIIISQs. (**b)** Percentage differences in the amino acid sequences between human FVIIISQ and other mammalian FVIIISQs. **(c, d)** Increases in FVIII activity (OSA and CSA) in the supernatant of HEK293 **(c)** or Huh-7 **(d)** cells transfected with plasmid. Values are presented as the mean ± SEM (*n* = 3). Significance was assessed using Student's *t*-tests.

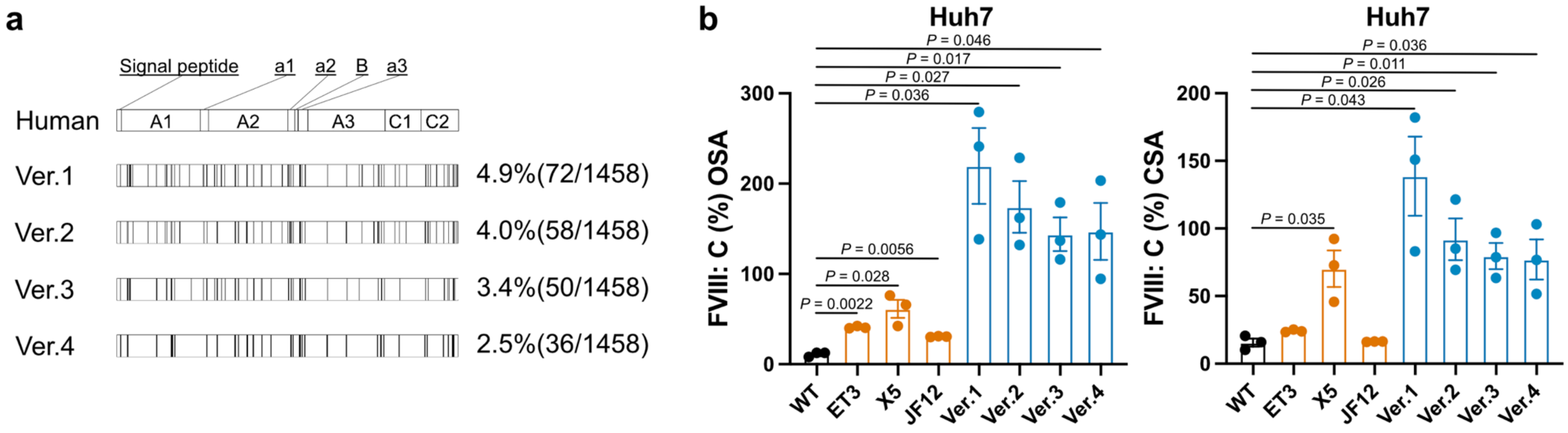

**Supplementary Fig. 2 | FVIII activity in the supernatant of Huh-7 cells transfected with plasmid harboring the engineered FVIIISQ.**
**(a)** Percentages and numbers of amino acid substitutions in the engineered FVIIISQs. (**b)** Increases in FVIII activity (OSA and CSA) in the supernatant of Huh-7 cells transfected with plasmid. Values are presented as the mean ± SEM (*n* = 3). Significance was assessed using Student's *t*-tests.

Ver, engineered FVIIIs in our study. ET3, X5, and JF12 are engineered FVIIISQs from the previous studies.

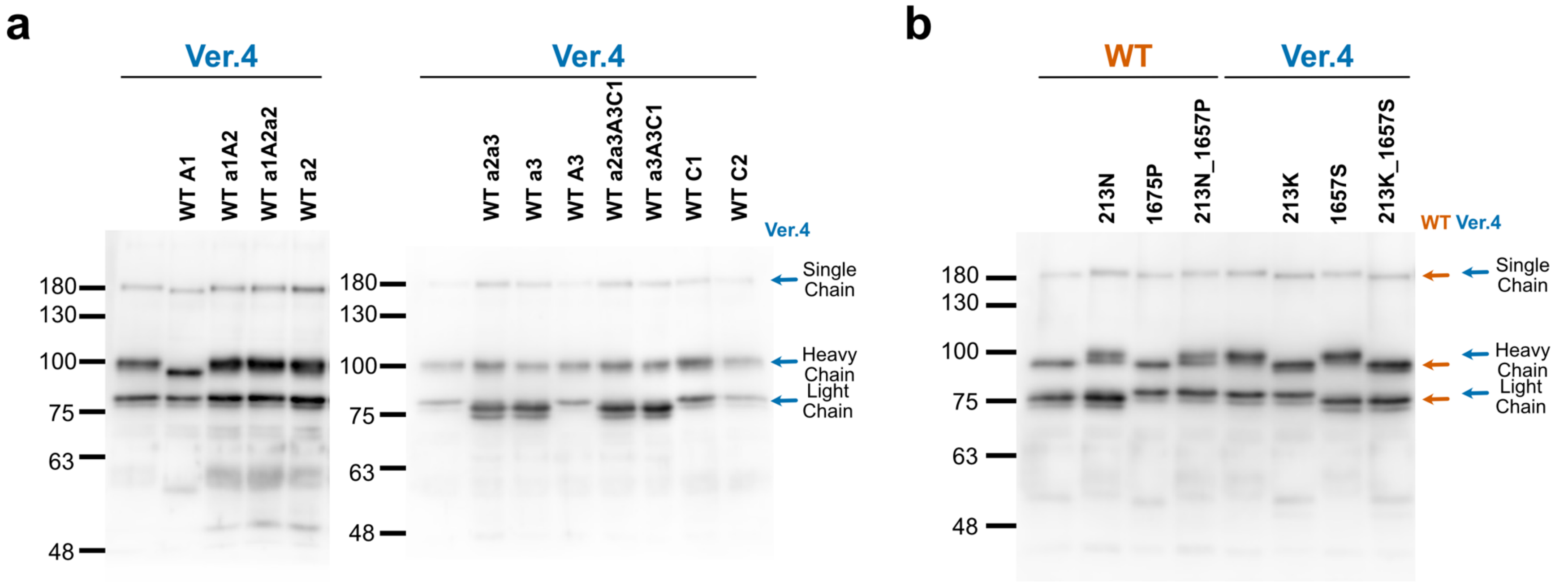

**Supplementary Fig. 3 | Identification of the amino acid substitutions in the engineered FVIIISQ(Ver.4) responsible for the increased molecular size.**

Immunoblotting analysis with anti-human FVIII antibody of supernatant from the Huh-7 cells transfected with FVIIISQ plasmids. **(a)** Immunoblot comparison of FVIIISQ(Ver.4) and FVIIISQ(Ver.4) with the indicated domains converted to those of WT. **(b)** Immunoblot comparison of FVIIISQ or FVIIISQ(Ver.4) without or with the indicated amino acid substitution(s). Representative immunoblotting images are shown. Arrows indicate the location of the single chain, heavy chain, and light chain of FVIIISQs.

WT, wild-type human B-domain-deleted factor VIII (FVIIISQ); Ver4, engineered FVIIISQ(Ver.4).

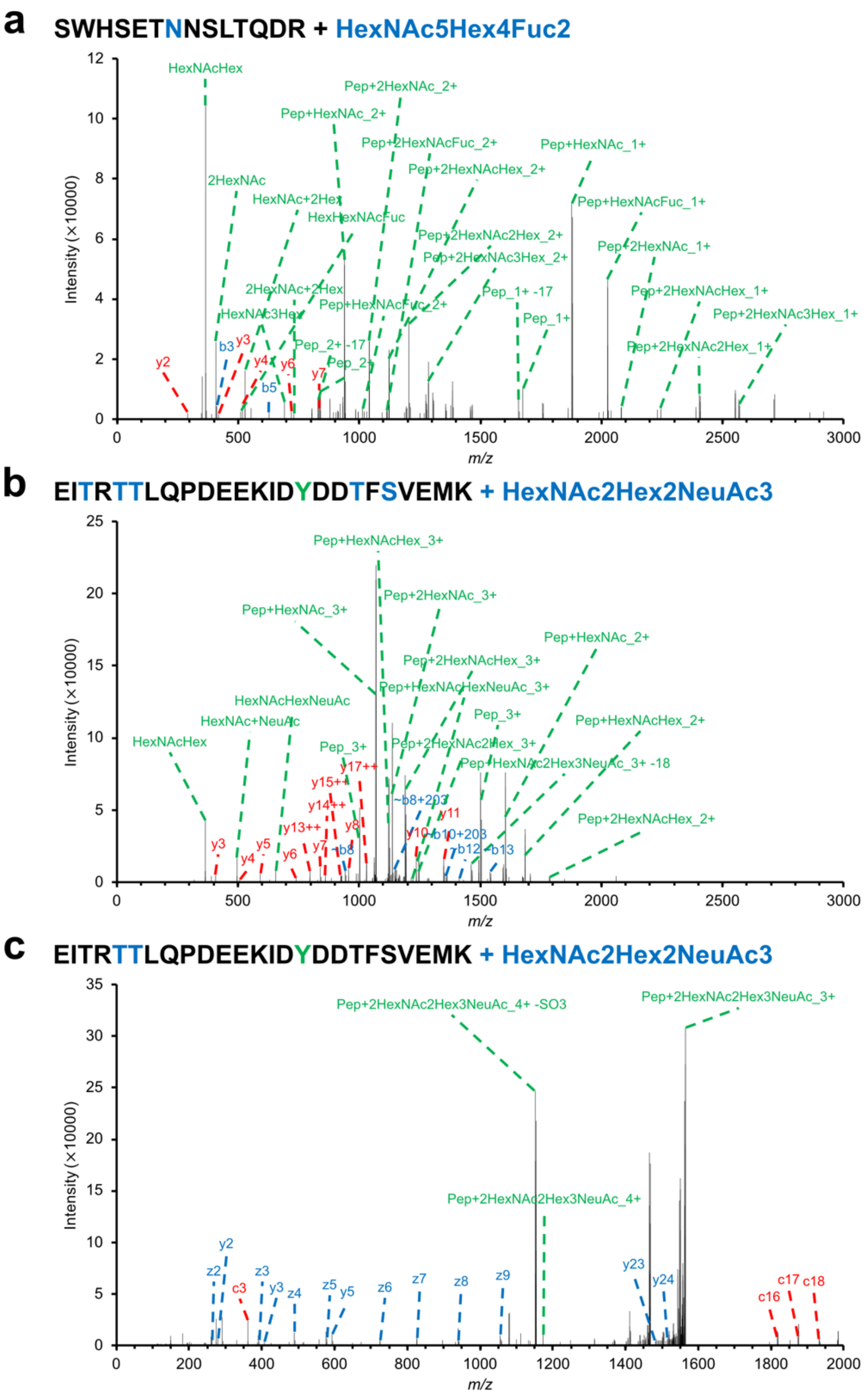

**Supplementary Fig. 4 | Glycosylation analysis of amino acid substitution sites that increased molecular size in the engineered FVIIISQ(Ver.4).**

**(a)** Higher-energy collisional dissociation mass spectrum of S207–R220 with HexNAc5Hex4Fuc2 identified in the heavy chain of FVIIISQ(Ver.4). **(b)** Higher-energy collisional dissociation mass spectra and **(c)** electron-transfer/higher-energy collisional dissociation mass spectra of E1649–K1673 with HexNAc2Hex2NeuAc3 identified in the light chain of FVIIISQ(Ver.4).

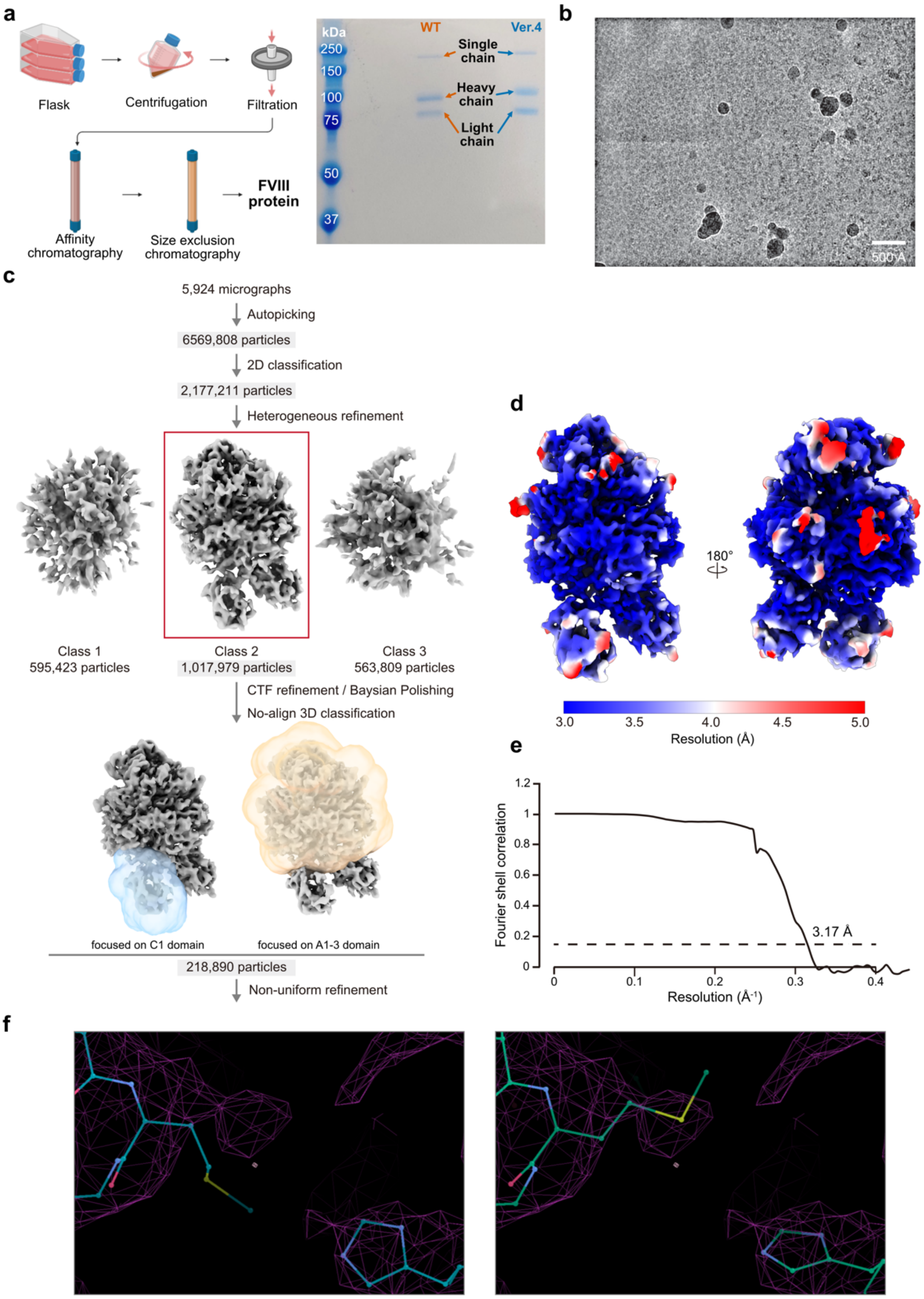

**Supplementary Fig. 5 | Structural data analysis and modeling.**
**(a)** Purification of FVIIISQ(Ver.4). **(b)** Representative cryo-electron microscopy (EM) images, recorded on a 300-kV Titan Krios microscope with a K3 camera. **(c)** Single-particle cryo-EM image processing workflows. (d) Final cryo-EM map colored by local resolution. (e) Fourier shell correlation (FSC) curves for the 3D reconstruction, with the gold-standard cut-off (FSC = 0.143) marked with a black dashed line. (f) Original (left) and refined (right) versions of the wild-type FVIII model (PDB:6MF2). The refined version is used for Fig. 4c,d.

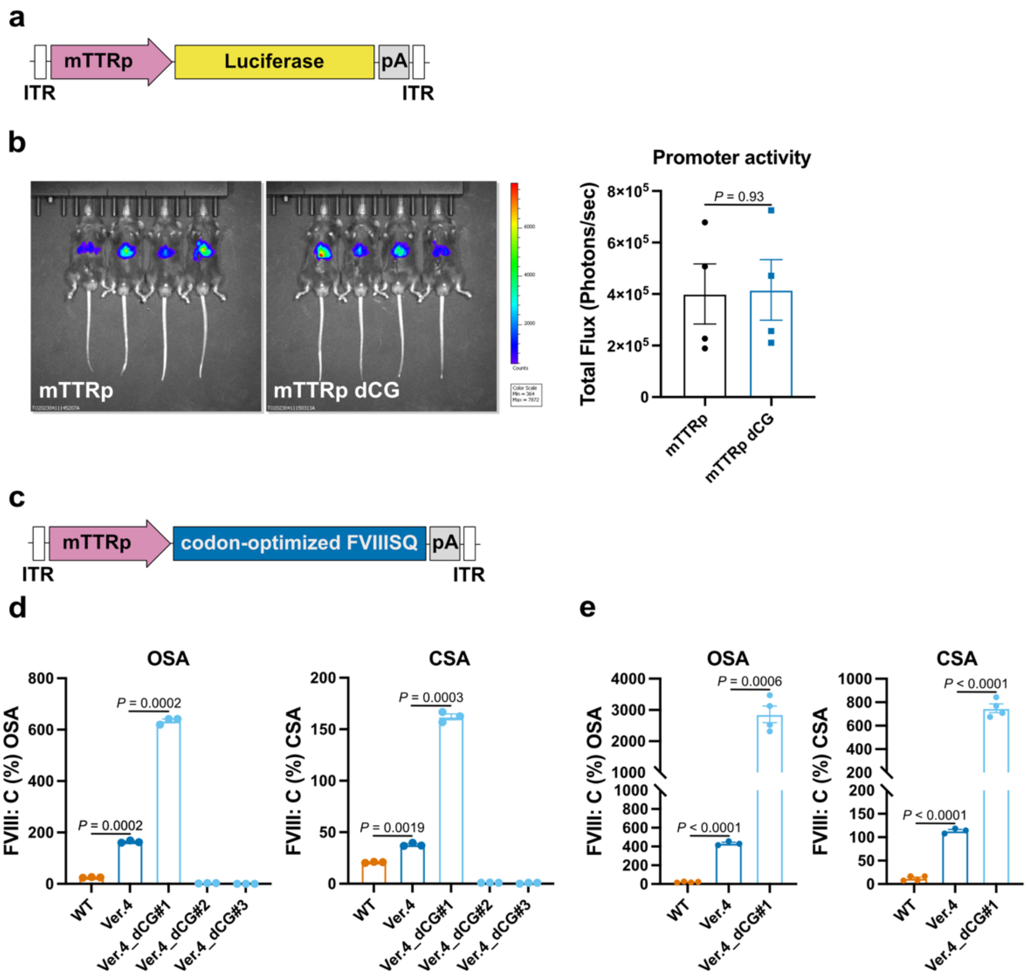

**Supplementary Fig. 6 | Optimization of the FVIIISQ(Ver.4) expression cassette to exclude CpG sequences.**

**(a)** Schematic presentation of the Adeno-associated virus (AAV) vector. The luciferase gene is driven by mouse transthyretin promoter (mTTRp). **(b)** C57BL/6 mice were treated with AAV8 vector harboring the luciferase gene driven by mTTRp or mTTRp without CpG sequences (mTTRp_dCG) ($1 \times 10^{10}$ vector genome [vg]). Representative IVIS imaging system data at 6 weeks after vector administration. Quantitative data are expressed as photon units (photons/s). Values are presented as the mean ± SEM ($n$ = 4). **(c)** Schematic presentation of the AAV vector. Each FVIIISQ gene is driven by mTTRp_dCG. **(d)** Increases in FVIII activity in the supernatants of Huh-7 cells transduced with AAV6 vector harboring the indicated FVIIISQ ($1 \times 10^{5}$ vg/cell). Values are presented as the mean ± SEM ($n$ = 3). **(e)** AAV8 vectors harboring the indicated FVIIISQ were intravenously administered into hemophilia A model mice ($1 \times 10^{11}$ vg/kg). The increases in plasma FVIII activity at 2 weeks after vector injection are shown. Values are presented as the mean ± SEM ($n$ = 4). Significance was assessed using Student's $t$-tests. WT, codon-optimized wild-type FVIIISQ; Ver.4: codon-optimized engineered FVIIISQ(Ver.4); Ver.4_dCG: codon-optimized engineered FVIIISQ(Ver.4) without CpG sequences. #1, #2, and #3 refer to the codon-optimized sequences from different companies.

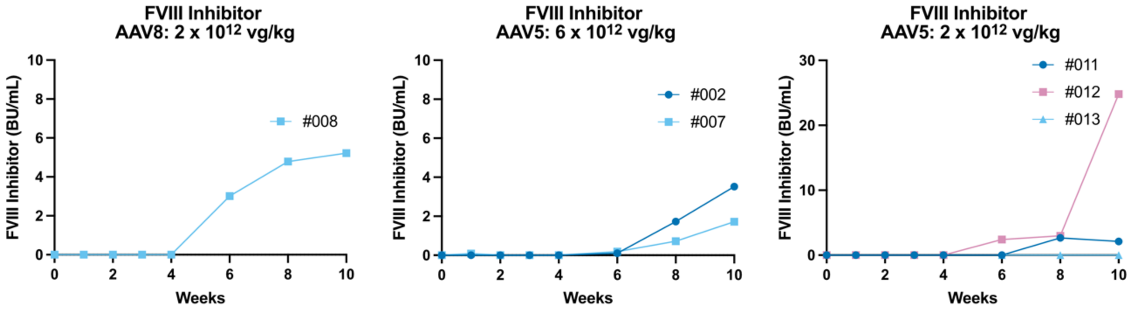

**Supplementary Fig. 7 | Emergence of neutralizing antibodies (inhibitors) against human FVIII in the serum of macaques treated with AAV vector.**

AAV8 or AAV5 harboring the codon-optimized engineered FVIIISQ(Ver.4) driven by the mTTR promoter without CpG sequences was administered to male macaques through a peripheral vein (AAV8: 2 × 10 vg/kg$^{12}$, #008; AAV5: 6 × $10^{12}$ vg/kg, #002, #007; AAV5: 2 × $10^{12}$ vg/kg, #011, #012, #013). The presence of neutralizing antibodies in the plasma was assessed using the Bethesda method. One Bethesda unit (BU/mL) refers to the inhibitor titer that inhibits 50% of human FVIII activity.

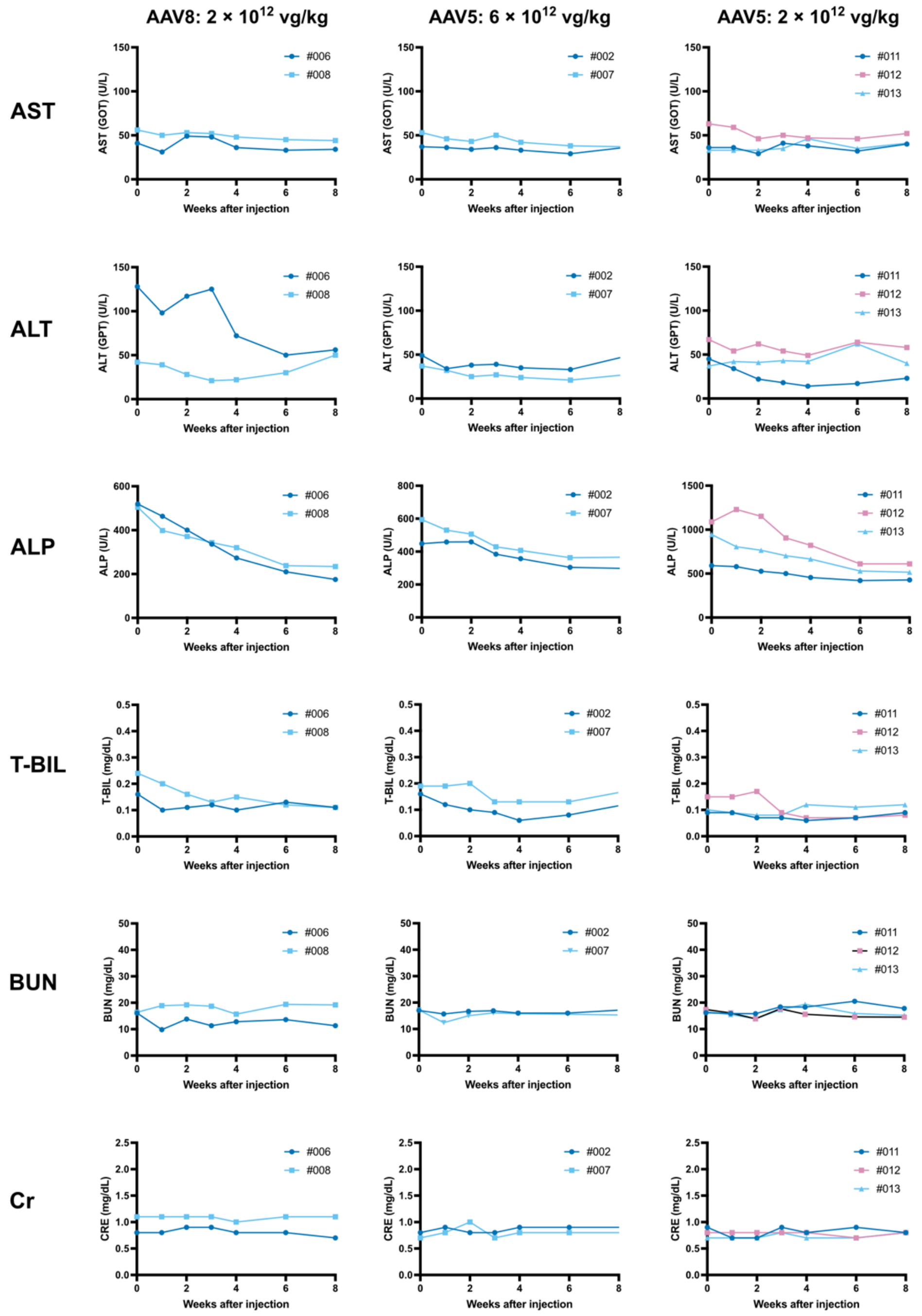

**Supplementary Fig. 8 | Changes in laboratory test parameters after AAV vector administration in macaques.**

AAV8 or AAV5 harboring the codon-optimized engineered FVIIISQ(Ver.4) driven by the mTTR promoter without CpG sequences was administered to male macaques through a peripheral vein (AAV8: 2 × 10 vg/kg$^{12}$, #008; AAV5: 6 × $10^{12}$ vg/kg, #002, #007; AAV5: 2 × $10^{12}$ vg/kg, #011, #012, #013). Changes in serum aspartate aminotransferase (AST), alanine aminotransferase (ALT), alkaline phosphatase (ALP), total bilirubin (T-BiL), blood urea nitrogen (BUN), and creatinine (Cr) were measured.

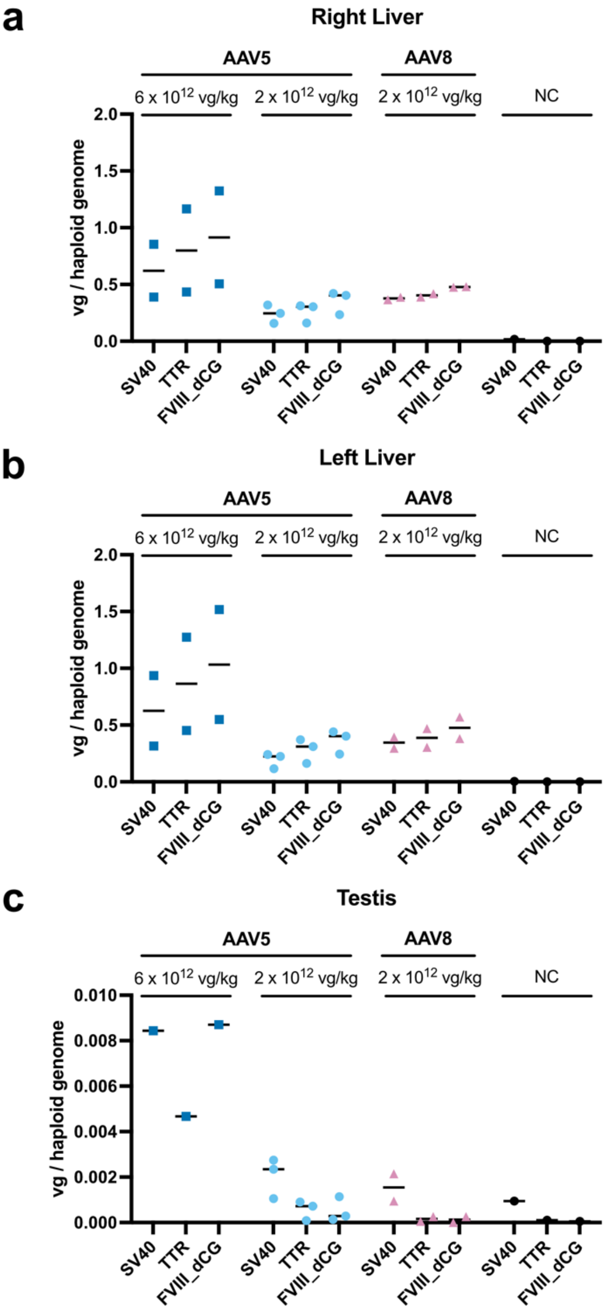

**Supplementary Fig. 9 | Digital PCR analysis of the numbers of transduced vector copies in the liver and testes of AAV vector-administered macaques.**

AAV8 or AAV5 harboring the codon-optimized engineered FVIIISQ(Ver.4) driven by the mTTR promoter without CpG sequences was administered to male macaques through a peripheral vein (AAV5: 6 × $10^{12}$ vg/kg, n=2; AAV5: 2 × $10^{12}$ vg/kg, n=3; AAV8: 2 × 10 vg/kg$^{12}$ n=2). AAV genome in the right **(a)** and left **(b)** liver and the testes (**c**), analyzed using digital PCR. The negative control (NC) was genomic DNA derived from untreated macaque liver tissue (n=1).

Fig. 2c

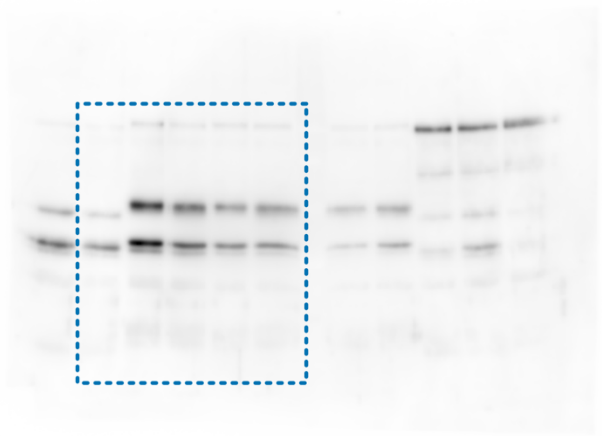

Fig. 2e FVIII

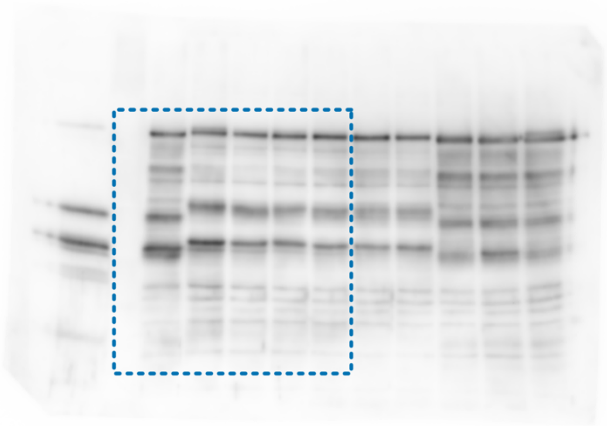

Fig. 2e β-actin

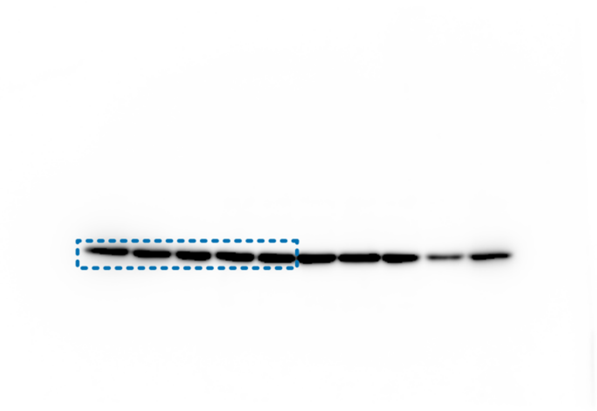

**Supplementary Fig. 10 | Original image data of Fig. 2**

Supplementary Fig.3a right

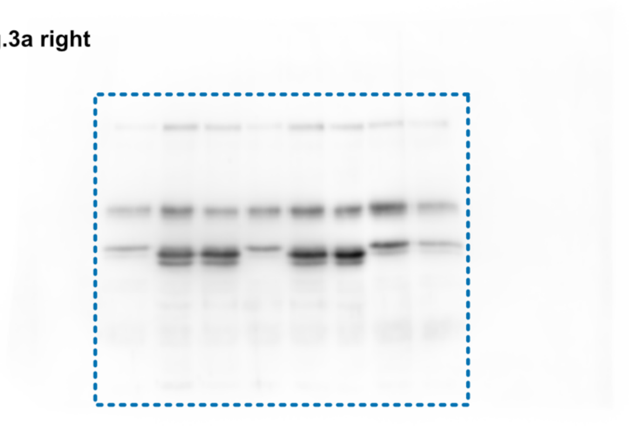

Supplementary Fig.3a left

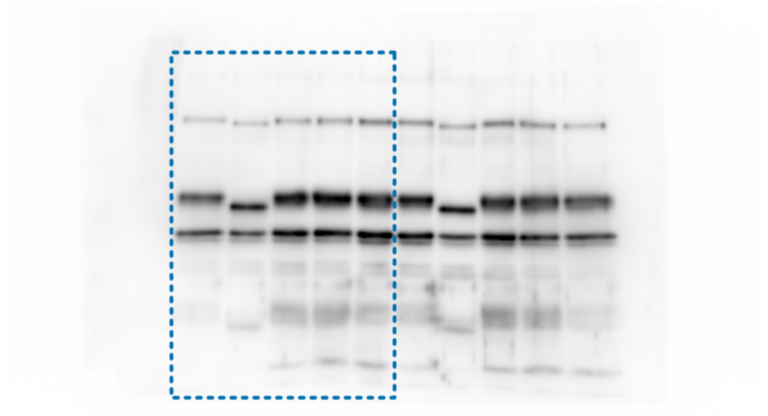

Supplementary Fig.3b

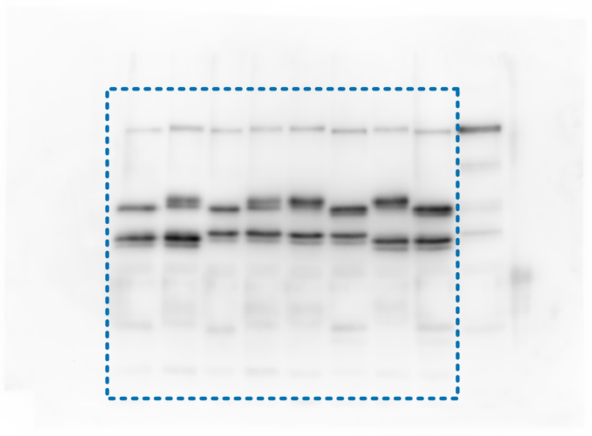

**Supplementary Fig. 11 | Original image data of Supplementary Fig. 3**

Supplementary Fig. 5a

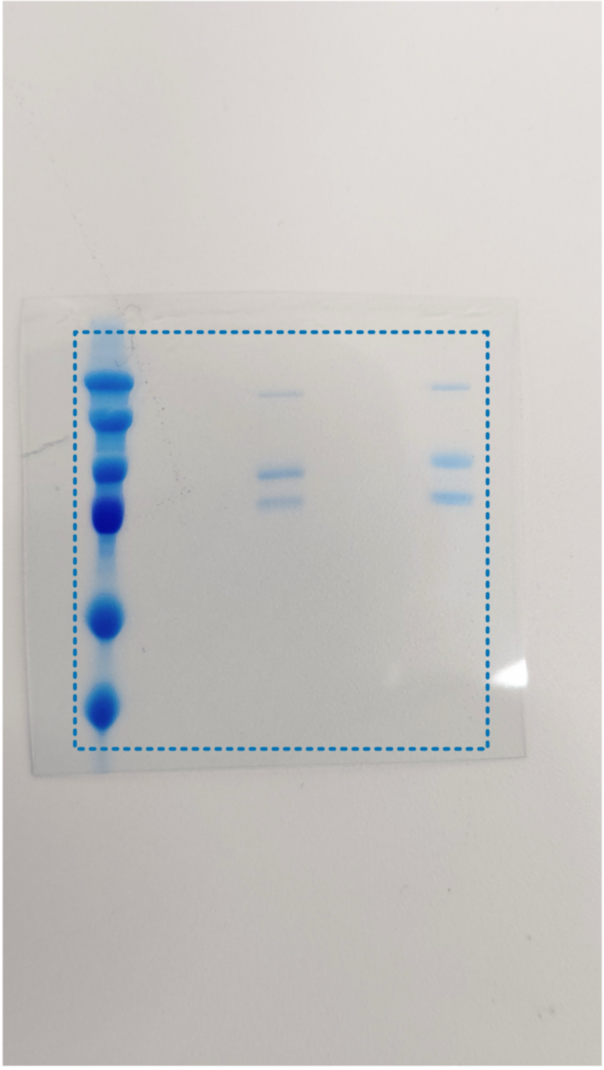

Supplementary Fig. 12 | Original image data of Supplementary Fig. 5

| | Ver.1 | Ver.2 | Ver.3 | Ver.4 |
|---|---|---|---|---|
| Signal | R-5P | R-5P | R-5P | R-5P |
| A1 | P25H | P25H | P25H | P25H |
| | A28T | A28T | A28T | A28T |
| | K36G | | K36G | |
| | F38L | | F38L | |
| | F40L | | F40L | |
| | L50V | L50V | | |
| | D115E | D115E | | |
| | L152P | L152P | L152P | L152P |
| | K188R | K188R | | |
| | K194E | | K194E | |
| | I196V | I196V | | |
| | K213N | K213N | K213N | K213N |
| | M217T | M217T | M217T | M217T |
| | W228Q | W228Q | W228Q | W228Q |
| | R250K | R250K | | |
| | L299F | | L299F | |
| a1 | T351Y | | T351Y | |
| | E354D | E354D | | |
| | P366S | | P366S | |
| | S367P | S367P | S367P | S367P |

| | Ver.1 | Ver.2 | Ver.3 | Ver.4 |
|---|---|---|---|---|
| A2 | L400P | | L400P | |
| | Q410L | Q410L | Q410L | Q410L |
| | M429V | | M429V | |
| | H444Y | | H444Y | |
| | Y487H | Y487H | Y487H | Y487H |
| | R489G | R489G | R489G | R489G |
| | F501M | F501M | F501M | F501M |
| | V537I | V537I | | |
| | M539L | M539L | M539L | M539L |
| | I566M | I566M | I566M | I566M |
| | A599D | | A599D | |
| | L603P | L603P | L603P | L603P |
| | I642V | I642V | I642V | I642V |
| | I689V | I689V | | |
| a2 | K713R | K713R | | |
| | T715I | | T715I | |
| | S722T | S722T | | |
| | S727P | S727P | S727P | S727P |
| | A736V | A736V | A736V | A736V |
| a3 | S1657P | S1657P | S763P | S1657P |
| | D1658E | D1658E | | |
| | Q1659E | Q1659E | Q1659E | Q1659E |
| | E1660D | E1660D | | |
| | E1661K | E1661K | E1661K | E1661K |
| | I1668F | I1668F | I1668F | I1668F |
| | D1681G | D1681G | D1681G | D1681G |

| | Ver.1 | Ver.2 | Ver.3 | Ver.4 |
|---|---|---|---|---|
| A3 | R1776K | R1776K | R1776K | R1776K |
| | V1857I | V1857I | | |
| | H1859R | H1859R | H1859R | H1859R |
| | F1912L | | F1912L | |
| | I1925V | I1925V | | |
| | L1975V | L1975V | | |
| | A1993V | A1993V | A1993V | A1993V |
| | V1998I | V1998I | | |
| | H2007Q | H2007Q | H2007Q | H2007Q |
| | N2019K | N2019K | N2019K | N2019K |
| C1 | E2066D | E2066D | | |
| | K2085M | K2085M | K2085M | K2085M |
| | T2114S | T2114S | | |
| C2 | F2196L | F2196L | F2196L | F2196L |
| | T2197S | T2197S | | |
| | K2207Q | K2207Q | K2207Q | K2207Q |
| | S2216T | S2216T | | |
| | V2243I | V2243I | | |
| | F2275L | F2275L | F2275L | F2275L |
| | F2290S | | F2290S | |
| | S2296A | S2296A | S2296A | S2296A |
| | V2314A | V2314A | V2314A | V2314A |
| | Q2316H | Q2316H | Q2316H | Q2316H |
| | M2321L | M2321L | M2321L | M2321L |
| | D2330Q | D2330Q | | |

**Supplementary Table 1**

| FVIII form | Specific activity (One stage assay) | Specific activity (Xa generation assay) |
|---|---|---|
| FVIIISQ WT | 1 | 1 |
| FVIIISQ(Ver.4) | 3.6 | 1.5 |

**Supplementary Table 2**

#### FVIIISQ-WT

| Allele Group | Allele | Epitope | EL Score | Predicted Affinity [nM] | Allele Frequency | T-Score |
|---|---|---|---|---|---|---|
| DRB3_4_5 | DRB3_0202 | YLLSKNNAI | 0.899712 | 169.81 | 34.3 | 0.202 |
| DRB3_4_5 | DRB4_0101 | LRMEVLGCE | 0.818214 | 212.26 | 41.8 | 0.197 |
| DRB3_4_5 | DRB4_0101 | MEVLGCEAQ | 0.924332 | 351.09 | 41.8 | 0.119 |
| DRB3_4_5 | DRB3_0301 | YLLSKNNAI | 0.819704 | 134.83 | 13 | 0.096 |
| DRB3_4_5 | DRB4_0101 | IHPQSWVHQ | 0.126854 | 436.34 | 41.8 | 0.096 |
| DRB1 | DRB1_0701 | YLLSKNNAI | 0.980888 | 153.58 | 13.5 | 0.088 |
| DRB3_4_5 | DRB4_0101 | WVHQIALRM | 0.010686 | 490.71 | 41.8 | 0.085 |
| DRB3_4_5 | DRB3_0101 | FDDDNSPSF | 0.907809 | 350.31 | 26.1 | 0.075 |
| DQ | HLA-DQA10401-DQB10402 | FATWSPSKA | 0.030681 | 204.88 | 12.8 | 0.062 |
| DRB3_4_5 | DRB5_0101 | IKTQGARQK | 0.406746 | 273.98 | 16 | 0.058 |
| DRB3_4_5 | DRB3_0101 | IDYDDTISV | 0.946334 | 469.95 | 26.1 | 0.056 |
| DRB1 | DRB1_0101 | YLLSKNNAI | 0.93575 | 98.97 | 5.4 | 0.055 |
| DQ | HLA-DQA10401-DQB10402 | IHGIKTQGA | 0.173954 | 239.42 | 12.8 | 0.053 |
| DRB1 | DRB1_0301 | IMSDKRNVI | 0.990062 | 260.96 | 13.7 | 0.052 |
| DRB3_4_5 | DRB5_0101 | FRNQASRPY | 0.1817 | 321.71 | 16 | 0.050 |
| DRB1 | DRB1_0101 | WVHQIALRM | 0.539068 | 109.77 | 5.4 | 0.049 |
| DRB1 | DRB1_1302 | YLLSKNNAI | 0.795101 | 170.92 | 7.7 | 0.045 |
| DRB3_4_5 | DRB3_0301 | LSKNNAIEP | 0.416955 | 289.57 | 13 | 0.045 |
| DRB3_4_5 | DRB5_0101 | FATWSPSKA | 0.21044 | 367.78 | 16 | 0.044 |
| DQ | HLA-DQA10102-DQB10602 | VHQIALRME | 0.631512 | 364.52 | 14.6 | 0.040 |
| DQ | HLA-DQA10401-DQB10402 | WSPSKARLH | 0.015011 | 325.63 | 12.8 | 0.039 |
| DP | HLA-DPA10201-DPB10101 | LRMEVLGCE | 0.140922 | 444.51 | 16 | 0.036 |
| DRB1 | DRB1_0101 | MEVLGCEAQ | 0.306461 | 154.15 | 5.4 | 0.035 |
| DRB1 | DRB1_0101 | VRPLYSRRL | 0.82351 | 161.09 | 5.4 | 0.034 |
| DQ | HLA-DQA10102-DQB10602 | IHGIKTQGA | 0.728759 | 437.95 | 14.6 | 0.033 |
| DRB3_4_5 | DRB5_0101 | WSPSKARLH | 0.013366 | 489.45 | 16 | 0.033 |
| DQ | HLA-DQA10401-DQB10402 | PSFIQIRSV | 0.068039 | 416.92 | 12.8 | 0.031 |
| DRB1 | DRB1_1501 | LRIHPQSWV | 0.116887 | 406.14 | 12.2 | 0.030 |
| DRB1 | DRB1_1501 | MVTFRNQAS | 0.940392 | 413.09 | 12.2 | 0.030 |
| DRB1 | DRB1_0701 | WVHQIALRM | 0.248702 | 470.7 | 13.5 | 0.029 |
| DQ | HLA-DQA10401-DQB10402 | FRNQASRPY | 0.011238 | 460.09 | 12.8 | 0.028 |
| DRB3_4_5 | DRB3_0301 | IMSDKRNVI | 0.52733 | 481.05 | 13 | 0.027 |
| DQ | HLA-DQA10401-DQB10402 | PYNIYPHGI | 0.004758 | 485.16 | 12.8 | 0.026 |
| DQ | HLA-DQA10401-DQB10402 | MVTFRNQAS | 0.038181 | 492.92 | 12.8 | 0.026 |
| DRB1 | DRB1_0101 | YEDISAYLL | 0.960955 | 237.23 | 5.4 | 0.023 |
| DRB1 | DRB1_1402 | YLLSKNNAI | 0.853206 | 246.79 | 5.6 | 0.023 |
| DRB1 | DRB1_0405 | FTPVVNSLD | 0.694466 | 283.84 | 6.2 | 0.022 |
| DRB1 | DRB1_0405 | YYSSFVNME | 0.351131 | 291.65 | 6.2 | 0.021 |
| DRB1 | DRB1_1402 | MEVLGCEAQ | 0.831141 | 306.87 | 5.6 | 0.018 |
| DRB1 | DRB1_1302 | IALRMEVLG | 0.613281 | 437.65 | 7.7 | 0.018 |
| DRB1 | DRB1_1301 | IHGIKTQGA | 0.699317 | 376.46 | 6.3 | 0.017 |
| DRB1 | DRB1_0405 | MEVLGCEAQ | 0.780612 | 377.07 | 6.2 | 0.016 |
| DRB1 | DRB1_0101 | FRNQASRPY | 0.041934 | 338.91 | 5.4 | 0.016 |
| DRB1 | DRB1_0101 | IHGIKTQGA | 0.190514 | 350.06 | 5.4 | 0.015 |
| DRB1 | DRB1_1301 | LRIHPQSWV | 0.122572 | 411.88 | 6.3 | 0.015 |
| DRB1 | DRB1_0901 | YLLSKNNAI | 0.918318 | 426.08 | 6.2 | 0.015 |
| DRB1 | DRB1_0404 | IHGIKTQGA | 0.894606 | 277.38 | 3.8 | 0.014 |
| DRB1 | DRB1_0401 | YLLSKNNAI | 0.909681 | 336 | 4.6 | 0.014 |
| DRB1 | DRB1_1301 | VTFRNQASR | 0.91812 | 461.14 | 6.3 | 0.014 |
| DRB1 | DRB1_1401 | IMSDKRNVI | 0.948357 | 496.3 | 6.7 | 0.013 |
| DRB1 | DRB1_0101 | FTNMFATWS | 0.019277 | 404.49 | 5.4 | 0.013 |
| DRB1 | DRB1_0101 | FATWSPSKA | 0.134829 | 405.14 | 5.4 | 0.013 |
| DRB1 | DRB1_0101 | VNSLDPPLL | 0.703564 | 414.03 | 5.4 | 0.013 |
| DRB1 | DRB1_1301 | IALRMEVLG | 0.721392 | 496.42 | 6.3 | 0.013 |
| DRB1 | DRB1_1402 | WVHQIALRM | 0.45164 | 479.1 | 5.6 | 0.012 |
| DRB1 | DRB1_0404 | MEVLGCEAQ | 0.914283 | 334.28 | 3.8 | 0.011 |
| DRB1 | DRB1_0101 | VKHLKDFPI | 0.795238 | 477.67 | 5.4 | 0.011 |
| DRB1 | DRB1_0407 | YLLSKNNAI | 0.95186 | 462.18 | 4.8 | 0.010 |
| DRB1 | DRB1_1104 | IIHGIKTQG | 0.931913 | 276.59 | 2.8 | 0.010 |
| DRB1 | DRB1_0404 | MVTFRNQAS | 0.942395 | 392.21 | 3.8 | 0.010 |
| DRB1 | DRB1_1104 | IHGIKTQGA | 0.659217 | 301.32 | 2.8 | 0.009 |
| DRB1 | DRB1_0402 | IHGIKTQGA | 0.951596 | 270.77 | 2.2 | 0.008 |
| DRB1 | DRB1_1601 | YLLSKNNAI | 0.962801 | 279.49 | 1.9 | 0.007 |
| DRB1 | DRB1_0402 | MVTFRNQAS | 0.979013 | 371.94 | 2.2 | 0.006 |
| DRB1 | DRB1_1102 | IHGIKTQGA | 0.699317 | 376.46 | 2.2 | 0.006 |
| DRB1 | DRB1_1104 | IALRMEVLG | 0.965629 | 488.98 | 2.8 | 0.006 |
| DRB1 | DRB1_1102 | LRIHPQSWV | 0.122572 | 411.88 | 2.2 | 0.005 |
| DRB1 | DRB1_1102 | VTFRNQASR | 0.91812 | 461.14 | 2.2 | 0.005 |
| DRB1 | DRB1_1102 | IALRMEVLG | 0.721392 | 496.42 | 2.2 | 0.004 |
| DRB1 | DRB1_1103 | IIHGIKTQG | 0.735967 | 238.3 | 0.5 | 0.002 |
| DRB1 | DRB1_1103 | IHGIKTQGA | 0.168348 | 304.48 | 0.5 | 0.002 |
| DRB1 | DRB1_1103 | LRIHPQSWV | 0.057985 | 430.79 | 0.5 | 0.001 |
| DRB1 | DRB1_1103 | LHLQGRSNA | 0.664788 | 462.53 | 0.5 | 0.001 |
| DRB1 | DRB1_1103 | VTFRNQASR | 0.334928 | 494.64 | 0.5 | 0.001 |
| DRB1 | DRB1_1304 | LRIHPQSWV | 0.147473 | 293.79 | 0.2 | 0.001 |
| DRB1 | DRB1_1304 | IALRMEVLG | 0.915764 | 464.31 | 0.2 | 0.000 |

#### FVIIISQ(Ver.4)

| Allele Group | Allele | Epitope | EL Score | Predicted Affinity [nM] | Allele Frequency | T-Score |
|---|---|---|---|---|---|---|
| DRB3_4_5 | DRB3_0202 | YLLSKNNVI | 0.941417 | 149.45 | 34.3 | 0.230 |
| DRB3_4_5 | DRB4_0101 | LEVLGCEAQ | 0.826491 | 258.66 | 41.8 | 0.162 |
| DRB1 | DRB1_0701 | YLLSKNNVI | 0.992951 | 110.54 | 13.5 | 0.122 |
| DRB3_4_5 | DRB4_0101 | IHPQSWAHH | 0.048149 | 371.79 | 41.8 | 0.112 |
| DRB3_4_5 | DRB4_0101 | LRLEVLGCE | 0.508221 | 374.97 | 41.8 | 0.111 |
| DRB1 | DRB1_0701 | WAHHIALRL | 0.61141 | 124.4 | 13.5 | 0.109 |
| DRB3_4_5 | DRB3_0101 | FDDDNSPPF | 0.929279 | 279.41 | 26.1 | 0.093 |
| DRB3_4_5 | DRB3_0301 | YLLSKNNVI | 0.872589 | 139.18 | 13 | 0.093 |
| DRB1 | DRB1_0101 | YLLSKNNVI | 0.980657 | 70.18 | 5.4 | 0.077 |
| DP | HLA-DPA10103-DPB10401 | WAHHIALRL | 0.18058 | 491.78 | 36.2 | 0.074 |
| DP | HLA-DPA10103-DPB10201 | WAHHIALRL | 0.382901 | 272.13 | 17.5 | 0.064 |
| DRB1 | DRB1_1302 | YLLSKNNVI | 0.917088 | 143.79 | 7.7 | 0.054 |
| DP | HLA-DPA10202-DPB10501 | LRLEVLGCE | 0.095617 | 407.83 | 21.7 | 0.053 |
| DQ | HLA-DQA10401-DQB10402 | FATWSPSQA | 0.027945 | 261.11 | 12.8 | 0.049 |
| DRB3_4_5 | DRB5_0101 | IMTQGARQK | 0.238823 | 329.23 | 16 | 0.049 |
| DP | HLA-DPA10201-DPB10101 | WAHHIALRL | 0.131159 | 334.13 | 16 | 0.048 |
| DP | HLA-DPA10202-DPB10501 | WAHHIALRL | 0.067749 | 466.81 | 21.7 | 0.046 |
| DQ | HLA-DQA10401-DQB10402 | IHGIMTQGA | 0.131305 | 294.31 | 12.8 | 0.043 |
| DP | HLA-DPA10103-DPB10201 | HIALRLEVL | 0.211502 | 404.25 | 17.5 | 0.043 |
| DRB1 | DRB1_0101 | WAHHIALRL | 0.376701 | 124.88 | 5.4 | 0.043 |
| DQ | HLA-DQA10102-DQB10602 | AHHIALRLE | 0.6227 | 359.97 | 14.6 | 0.041 |
| DQ | HLA-DQA10401-DQB10402 | ALRLEVLGC | 0.135279 | 316.9 | 12.8 | 0.040 |
| DQ | HLA-DQA10102-DQB10602 | IHGIMTQGA | 0.546889 | 392.69 | 14.6 | 0.037 |
| DRB3_4_5 | DRB5_0101 | YLLSKNNVI | 0.847734 | 449.8 | 16 | 0.036 |
| DRB1 | DRB1_1501 | MVTFKNQAS | 0.962975 | 358.94 | 12.2 | 0.034 |
| DQ | HLA-DQA10401-DQB10402 | PQSWAHHIA | 0.041024 | 380.3 | 12.8 | 0.034 |
| DRB3_4_5 | DRB3_0301 | LLVCRTNTL | 0.168372 | 395.34 | 13 | 0.033 |
| DRB1 | DRB1_0101 | YEDIPAYLL | 0.967337 | 168.57 | 5.4 | 0.032 |
| DRB1 | DRB1_0901 | WAHHIALRL | 0.484871 | 218.52 | 6.2 | 0.028 |
| DRB1 | DRB1_1501 | IALRLEVLG | 0.685369 | 441.78 | 12.2 | 0.028 |
| DRB1 | DRB1_0405 | FTPVVNALD | 0.722685 | 226.82 | 6.2 | 0.027 |
| DQ | HLA-DQA10401-DQB10402 | WSPSQARLH | 0.012964 | 474.07 | 12.8 | 0.027 |
| DRB1 | DRB1_0101 | LEVLGCEAQ | 0.332378 | 203.78 | 5.4 | 0.026 |
| DRB3_4_5 | DRB3_0301 | CRTNTLNPA | 0.226376 | 497.49 | 13 | 0.026 |
| DRB1 | DRB1_0101 | VRPLHSGRL | 0.804675 | 215.53 | 5.4 | 0.025 |
| DRB1 | DRB1_1501 | LRIHPQSWA | 0.09 | 493.68 | 12.2 | 0.025 |
| DRB1 | DRB1_1402 | YLLSKNNVI | 0.936361 | 227.77 | 5.6 | 0.025 |
| DRB1 | DRB1_1302 | IALRLEVLG | 0.656145 | 346.15 | 7.7 | 0.022 |
| DRB3_4_5 | DRB5_0102 | YLLSKNNVI | 0.855229 | 478.91 | 9.8 | 0.020 |
| DRB3_4_5 | DRB5_0102 | LEVLGCEAQ | 0.564419 | 490.08 | 9.8 | 0.020 |
| DRB1 | DRB1_0405 | YYSSFVNLE | 0.448289 | 311.17 | 6.2 | 0.020 |
| DRB1 | DRB1_1402 | LEVLGCEAQ | 0.848385 | 281.62 | 5.6 | 0.020 |
| DRB1 | DRB1_1302 | WAHHIALRL | 0.365728 | 426.61 | 7.7 | 0.018 |
| DRB1 | DRB1_1301 | IALRLEVLG | 0.741927 | 366.05 | 6.3 | 0.017 |
| DRB1 | DRB1_0901 | YLLSKNNVI | 0.970959 | 366.71 | 6.2 | 0.017 |
| DRB1 | DRB1_0101 | VNALDPPLL | 0.810127 | 324.96 | 5.4 | 0.017 |
| DRB1 | DRB1_1302 | FLQNGKVKV | 0.143619 | 477.51 | 7.7 | 0.016 |
| DRB1 | DRB1_1401 | HHIALRLEV | 0.401271 | 437.06 | 6.7 | 0.015 |
| DRB1 | DRB1_1301 | VTFKNQASR | 0.941617 | 414.26 | 6.3 | 0.015 |
| DRB1 | DRB1_0405 | LEVLGCEAQ | 0.790474 | 437.4 | 6.2 | 0.014 |
| DRB1 | DRB1_1301 | MTQGARQKF | 0.082894 | 447.94 | 6.3 | 0.014 |
| DRB1 | DRB1_1301 | MIIHGIMTQ | 0.350543 | 450.78 | 6.3 | 0.014 |
| DRB1 | DRB1_0101 | YKSLYLNNG | 0.506236 | 403.8 | 5.4 | 0.013 |
| DRB1 | DRB1_0404 | IHGIMTQGA | 0.754839 | 298.73 | 3.8 | 0.013 |
| DRB1 | DRB1_0101 | IHGIMTQGA | 0.105953 | 425.17 | 5.4 | 0.013 |
| DRB1 | DRB1_0901 | FFSGYTFKH | 0.014214 | 488.25 | 6.2 | 0.013 |
| DRB1 | DRB1_1402 | WAHHIALRL | 0.528103 | 448.09 | 5.6 | 0.012 |
| DRB1 | DRB1_0401 | YLLSKNNVI | 0.927365 | 394.98 | 4.6 | 0.012 |
| DRB1 | DRB1_0404 | LEVLGCEAQ | 0.974556 | 359.46 | 3.8 | 0.011 |
| DRB1 | DRB1_0407 | YLLSKNNVI | 0.955054 | 469.97 | 4.8 | 0.010 |
| DRB1 | DRB1_0802 | IALRLEVLG | 0.828201 | 488.84 | 4.9 | 0.010 |
| DRB1 | DRB1_0404 | MVTFKNQAS | 0.959719 | 388.04 | 3.8 | 0.010 |
| DRB1 | DRB1_1601 | YLLSKNNVI | 0.985762 | 219.7 | 1.9 | 0.009 |
| DRB1 | DRB1_1104 | IALRLEVLG | 0.968551 | 415.37 | 2.8 | 0.007 |
| DRB1 | DRB1_0402 | MVTFKNQAS | 0.98371 | 352.43 | 2.2 | 0.006 |
| DRB1 | DRB1_0402 | IHGIMTQGA | 0.801901 | 357.39 | 2.2 | 0.006 |
| DRB1 | DRB1_1102 | IALRLEVLG | 0.741927 | 366.05 | 2.2 | 0.006 |
| DRB1 | DRB1_1102 | VTFKNQASR | 0.941617 | 414.26 | 2.2 | 0.005 |
| DRB1 | DRB1_1102 | MTQGARQKF | 0.082894 | 447.94 | 2.2 | 0.005 |
| DRB1 | DRB1_1102 | MIIHGIMTQ | 0.350543 | 450.78 | 2.2 | 0.005 |
| DRB1 | DRB1_1601 | WAHHIALRL | 0.350191 | 475.55 | 1.9 | 0.004 |
| DRB1 | DRB1_1103 | LRIHPQSWA | 0.129892 | 357.38 | 0.5 | 0.001 |
| DRB1 | DRB1_1103 | MTQGARQKF | 0.040543 | 449.03 | 0.5 | 0.001 |
| DRB1 | DRB1_1103 | IHGIMTQGA | 0.277013 | 466.21 | 0.5 | 0.001 |
| DRB1 | DRB1_1103 | IMVTFKNQA | 0.895598 | 483.49 | 0.5 | 0.001 |
| DRB1 | DRB1_1103 | VTFKNQASR | 0.472362 | 492.67 | 0.5 | 0.001 |
| DRB1 | DRB1_1304 | IALRLEVLG | 0.752197 | 376.47 | 0.2 | 0.001 |
| DRB1 | DRB1_1304 | MTQGARQKF | 0.096768 | 425.62 | 0.2 | 0.000 |

**Supplementary Table 3**

| Domain | FVIIISQ(WT) | Allele counts | FVIIISQ(Ver.4) | Allele counts | Increased number of antigenic epitopes (Ver. 4-WT) |
|---|---|---|---|---|---|
| a2 | YLLSKNNAI | 10 | YLLSKNNVI | 12 | 2 |
| C1 | IHGIKTQGA | 9 | IHGIMTQGA | 6 | -3 |
| C2 | MEVLGCEAQ | 5 | LEVLGCEAQ | 6 | 1 |
| C2 | LRIHPQSWV | 5 | LRIHPQSWA | 2 | -3 |
| C2 | IALRMEVLG | 5 | IALRLEVLG | 7 | 2 |
| C2 | WVHQIALRM | 4 | WAHHIALRL | 10 | 6 |
| A3 | MVTFRNQAS | 4 | MVTFKNQAS | 3 | -1 |
| C2 | FATWSPSKA | 3 | FATWSPSQA | 1 | -2 |
| A2 | IMSDKRNVI | 3 | | 0 | -3 |
| A3 | FRNQASRPY | 3 | | 0 | -3 |
| A3 | VTFRNQASR | 3 | VTFKNQASR | 3 | 0 |
| C2 | LRMEVLGCE | 2 | LRLEVLGCE | 2 | 0 |
| C2 | WSPSKARLH | 2 | WSPSQARLH | 1 | -1 |
| C1 | IIHGIKTQG | 2 | | 0 | -2 |
| C2 | IHPQSWVHQ | 1 | IHPQSWAHH | 1 | 0 |
| a1 | FDDDNSPSF | 1 | FDDDNSPPF | 1 | 0 |
| C1 | IKTQGARQK | 1 | IMTQGARQK | 1 | 0 |
| a3 | IDYDDTISV | 1 | | 0 | -1 |
| a2 | LSKNNAIEP | 1 | | 0 | -1 |
| C2 | VHQIALRME | 1 | AHHIALRLE | 1 | 0 |
| A2 | VRPLYSRRL | 1 | VRPLHSGRL | 1 | 0 |
| a1A2 | PSFIQIRSV | 1 | | 0 | -1 |
| A2 | PYNIYPHGI ** | 1 | | 0 | -1 |
| a2 | YEDISAYLL | 1 | YEDIPAYLL | 1 | 0 |
| C2 | FTPVVNSLD | 1 | FTPVVNALD | 1 | 0 |
| A2 | YYSSFVNME | 1 | YYSSFVNLE | 1 | 0 |
| C2 | FTNMFATWS | 1 | | 0 | -1 |
| C2 | VNSLDPPLL | 1 | VNALDPPLL | 1 | 0 |
| A2 | VKHLKDFPI | 1 | | 0 | -1 |
| C2 | LHLQGRSNA ** | 1 | | 0 | -1 |
| C1 | | | MTQGARQKF | 4 | 4 |
| C1 | | | MIIHGIMTQ | 2 | 2 |
| C2 | | | HIALRLEVL | 1 | 1 |
| C2 | | | ALRLEVLGC | 1 | 1 |
| C2 | | | PQSWAHHIA | 1 | 1 |
| A3 | | | LLVCRTNTL | 1 | 1 |
| A3 | | | CRTNTLNPA | 1 | 1 |
| C2 | | | FLQNGKVKV | 1 | 1 |
| C2 | | | HHIALRLEV | 1 | 1 |
| A2 | | | YKSLYLNNG | 1 | 1 |
| A2 | | | FFSGYTFKH * | 1 | 1 |
| A3 | | | IMVTFKNQA | 1 | 1 |

* Addition of antigenic epitope in Ver.4 by surrounding substitution sites.
** Reduction of antigenic epitope in Ver.4 by surrounding substitution sites.

**Supplementary Table 4**

| Epitope Sequence | Allele counts | T-score | | | |
|---|---|---|---|---|---|
| **M**TQGARQKF | 4 | 0.014064384 | 0.004911372 | 0.001113511 | 0.000469903 |
| MIIHGI**M**TQ | 2 | 0.013975775 | 0.004880429 | | |
| **H**IALR**L**EVL | 1 | 0.043290043 | | | |
| ALR**L**EVLGC | 1 | 0.040391291 | | | |
| PQSW**A**H**H**IA | 1 | 0.033657639 | | | |
| LLVC**R**TNTL | 1 | 0.032883088 | | | |
| C**R**TNTLNPA | 1 | 0.026131179 | | | |
| F**L**QNGKVKV | 1 | 0.016125317 | | | |
| H**H**IALR**L**EV | 1 | 0.015329703 | | | |
| YKS**L**YLNNG | 1 | 0.013372957 | | | |
| FFSGYTFKH * | 1 | 0.012698413 | | | |
| IMVTF**K**NQA | 1 | 0.001034148 | | | |

**Substitution sites in FVIIISQ(Ver.4)**

* Addition of antigenic epitope by surrounding substitution sites.

**Supplementary Table 5**

| | |
|---|---|
| For titration of AAV vectors and qPCR/dPCR of vg in genomic DNA | |
| SV40_F | 5' -GCAATAGCATCACAAATTTCAC-3′ |
| SV40_R | 5' -GATCCAGACATGATAAGATACATTG-3′ |
| SV40 Probe | 5' -TCACTGCATTCTAGTTGTGGTTTGTCCA-3′ |
| For qPCR of hFVIIISQ-CO in mRNA | |
| hFVIIISQqPCR_F | 5'- GTGAAGCCCAACGAGACTAA -3' |
| hFVIIISQqPCR_R | 5'- AGGTCCACATCGGAGAAGTA -3' |
| hFVIIISQqPCR_Probe | 5'- CCACAAAGGACGAGTTCGACTGCA -3' |
| For qPCR of hFVIIISQ-COΔCG in mRNA and dPCR of vg in genomic DNA | |
| hFVIIISQ_ΔCG_qPCR_F | 5'- TGGAGAGACTGTGGGACTATG -3' |
| hFVIIISQ_ΔCG_qPCR_R | 5'- GGAACACCACCTTCTTGAACT -3' |
| hFVIIISQ_ΔCG_qPCR_Probe | 5'- ATGTGCTGAGAAACAGAGCCCAGT -3' |
| For dPCR of vg in genomic DNA | |
| mTtrp_qPCR_F | 5'- CCATCTTACTCAACATCCTCCC -3' |
| mTtrp_qPCR_R | 5'- CAGCCTGGGTTGGAAGG -3' |
| mTtrp_qPCR_Probe | 5'- ATCCCTGCCAAGCTGACTCCAAA -3' |

**Supplementary Table 6**

| IDs | Number | Age | BW (kg) | Administered AAV serotype | Dose | NAbs | |
|---|---|---|---|---|---|---|---|
| | | | | | | AAV8 | AAV5 |
| 1521802001 | #002 | 5 | 4.1 | AAV5 | $6 \times 10^{12}$ vg/kg | + | – |
| 1521711049 | #006 | 6 | 4.3 | AAV8 | $1 \times 10^{12}$ vg/kg | – | NA |
| 1521712060 | #007 | 6 | 3.4 | AAV5 | $6 \times 10^{12}$ vg/kg | + | – |
| 1521710043 | #008 | 6 | 3.9 | AAV8 | $1 \times 10^{12}$ vg/kg | – | NA |
| 1422006056 | #011 | 3 | 2.9 | AAV5 | $2 \times 10^{12}$ vg/kg | – | – |
| 1421909106 | #012 | 4 | 3.3 | AAV5 | $2 \times 10^{12}$ vg/kg | – | – |
| 1421707052 | #013 | 6 | 4 | AAV5 | $2 \times 10^{12}$ vg/kg | – | – |

**Supplementary Table 7**
